# Hot and sick: impacts of warming and oomycete parasite infection on endemic dominant zooplankter of Lake Baikal

**DOI:** 10.1101/711655

**Authors:** Ted Ozersky, Teofil Nakov, Stephanie E. Hampton, Nicholas L. Rodenhouse, Kirill Shchapov, Kara H. Woo, Katie Wright, Helena V. Pislegina, Lyubov R. Izmest’eva, Eugene A. Silow, Maxim A. Timofeev, Marianne V. Moore

**Affiliations:** Large Lakes Observatory and Department of Biology, University of Minnesota Duluth, Duluth, Minnesota, USA.; Department of Biological Sciences, University of Arkansas, Fayetteville, Arkansas, USA.; School of the Environment, Washington State University, Pullman, Washington, USA.; Department of Biological Sciences, Wellesley College, Wellesley, Massachusetts, USA.; Sage Bionetworks, Seattle, Washington, USA.; Department of Biological Sciences, Wellesley College, Wellesley, Massachusetts, USA. Current address: Museum of Science, Boston, Massachusetts, USA.; Institute of Biology, Irkutsk State University, Irkutsk, Russia.

**Keywords:** Climate change, zooplankton, parasites, Lake Baikal, *Epischura baikalensis*, Oomycota, *Saprolegnia*, Diel Vertical Migration

## Abstract

Climate warming impacts ecosystems through multiple interacting pathways, including via direct thermal responses of individual taxa and the combined responses of closely interacting species. In this study we examined how warming and infection by an oomycete parasite affect the dominant zooplankter of Russia’s Lake Baikal, the endemic cold-adapted stenotherm *Epischura baikalensis* (Copepoda). We used a combination of laboratory experiments, long-term monitoring data and population modeling. Experiments showed large thermal mismatch between host and parasite, with strong negative effects of warm temperatures on *E. baikalensis* survival and reproduction and a negative synergistic effect of *Saprolegnia* infection. However, *Saprolegnia* infection had an unexpected positive effect on *E. baikalensis* reproductive output, which may be consistent with fecundity compensation by infected females. Long-term monitoring data showed that *Saprolegnia* infections were most common during the warmest periods of the year and that infected individuals tended to accumulate in deep water. Population models, parameterized with experimental and literature data, correctly predicted the timing of *Saprolegnia* epizootics, but overestimated the negative effect of warming on *E.baikalensis* populations. Models suggest that diel vertical migration may allow *E. baikalensis* to escape the negative effects of increasing temperatures and parasitism and enable *E. baikalensis* to persist as Lake Baikal warms. Our results contribute to understanding of how multiple interacting stressors affect warming pelagic ecosystems of cold lakes and oceans and show that the population-level consequences of thermal mismatch between hosts and parasites can vary seasonally, interannual and spatially.

## Introduction

The ecological effects of climate warming are manifested through multiple interacting physical, chemical and biological mechanisms (Adrian et al. 2009; Larsen et al. 2011; Zarnetske et al. 2012; O’Reilly et al. 2015; Kraemer et al. 2017). Among these mechanisms are the thermal tolerances of not only individual species but also the combined temperature responses of closely interacting taxa. The ecological significance of combined temperature responses has been shown for mutualistic, competitive and exploitative interactions (Kordas et al. 2011; Rafferty et al. 2015; Altizer et al. 2013; Cohen et al. 2017; Gehman et al. 2018). The effect of climate on these interactions is often idiosyncratic and affected by the specific life-histories of interacting organisms (Altizer et al. 2013; Warren and Bradford 2014; Cohen et al. 2017; Gehman et al. 2018) and by aspects of the abiotic environment, such as habitat spatial structure and the presence of refugia (Decaestecker et al. 2002; Araújo and Luoto 2007; Duffy 2007; Schweiger et al. 2008; Penczykowski et al. 2011). This complexity makes it challenging to predict how specific systems will respond to future climate changes and requires system-specific knowledge.

Parasites and diseases of phytoplankton and zooplankton provide examples of such complex interactions. They are common in lakes and oceans and can strongly affect the populations of the interacting species as well as ecosystem processes (Burns 1989; Ebert 2005; Duffy 2007; Ibelings et al. 2011; Valois and Poulin 2015; Valois and Burns 2016). Most of the work on freshwater parasites of zooplankton has involved *Daphnia* species and their bacterial and fungal parasites in small lakes and ponds (e.g., Ebert 2005, Duffy 2007; Wolinska et al. 2008; Hall et al. 2010; Duffy and Hunsberger 2018). Freshwater copepods are also affected by various parasites (Burns 1989; Miao and Nauwerck 1999; Valois and Burns 2016), but much less is known about their host-parasite interactions. While *Daphnia* often dominate zooplankton communities in small, warm and productive lakes and ponds, copepods are more important members of the pelagic plankton in large and cold lakes as well as in the world’s oceans (Gallienne and Robins 2001; Barbiero et al. 2019; Moore et al. 2019). Large lakes contain a disproportionate amount of global surface fresh water and support important ecosystem services, including valuable commercial fisheries. It is thus important to improve understanding of how temperature changes affect freshwater copepods through both direct effects and their interactions with parasites.

The pelagic food web of Russia’s Lake Baikal, the world’s oldest, most voluminous and most biologically diverse lake (Kozhov 1963; Moore et al. 2009), is dominated by a single species of zooplankton, the endemic calanoid copepod *Epischura baikalensis* Sars. *E. baikalensis* often comprises more than 90% of the pelagic zooplankton biomass in the cold waters of open Lake Baikal throughout the year but is conspicuously absent from the lake’s shallow bays during summer months, where cosmopolitan species of cladocerans, cyclopoids and calanoids are abundant (Kozhov 1963; Bowman 2014). Field data-based correlations and limited laboratory research have shown that *E. baikalensis* is a cold-adapted stenotherm that does not tolerate temperatures above ca. 15°C (Kozhov 1963; Afanasyeva 1977). Given recent warming trends in Lake Baikal and other large lakes (Shimaraev and Domysheva 2013, O’Reilly et al. 2015), there is concern that changes in the abundance of key species such as *E. baikalensis* may greatly alter lake food webs (Hampton et al. 2008; Moore et al. 2009). Warming temperatures may also affect copepods such as *E. baikalensis* through the negative impacts of parasites. As early as 1926, Lake Baikal researchers documented occasional mass epizootics of a fungal parasite, putatively identified as *Saprolegnia* sp. (Oomycota), that could impact almost the entire population of *E. baikalensis*, leading to large reductions in its abundance (Afanasyeva 1977). These epizootics were reported to occur during the summer months of particularly warm years and were believed to be at least partially temperature-related (Kozhov 1963; Afanasyeva 1977). Despite the potentially important effect of this parasite on *E. baikalensis*, no published detailed experimental or field studies have been conducted on the interactions between *E. baikalensis* and *Saprolegnia*.

The main objectives of this study were to assess the effects of temperature and parasite infection on an endemic dominant species in one of the world’s largest lakes and, more generally, to contribute to improved understanding of the impacts of climate change on the ecology of zooplankton in cold pelagic ecosystems. To achieve these objectives, we used lab experiments to assess the individual and combined responses of *E. baikalensis* and their oomycete parasite to a range of environmentally-relevant temperatures. Results of experiments were used to parameterize a population model that examined how different realistic temperature scenarios may affect *E. baikalensis* in open Lake Baikal and some of its warm bays and whether diel vertical migration (DVM) can offer *E. baikalensis* a thermal refuge during warm summer months. Long-term monitoring data were used to assess whether our population models produced realistic patterns of *E. baikalensis* abundance and parasite infection. We tested 5 specific hypotheses: H1) survival and reproduction of *E. baikalensis* will decrease with increasing temperatures; H2) prevalence and virulence of *Saprolegnia* will increase with increasing temperatures; H3) warmer temperatures and *Saprolegnia* exposure will interact synergistically, leading to larger reduction in *E. baikalensis* fitness than either factor alone; H4) *E. baikalensis* abundance will be lower under warmer conditions and in shallow, warm regions of Lake Baikal as revealed by long-term monitoring data and predicted by population models; H5) Population models will show that DVM provides a refuge for *E. baikalensis* from warm temperatures and parasitism.

## Methods

### Study system

#### Lake Baikal

Lake Baikal is located in Central Siberia and spans 600 km along its longest axis (Fig. S1). Due to the cold regional climate, Lake Baikal is frozen for up to 5 months of the year (January–May). The lake’s upper 300-m water layer is dimictic with a relatively short stratified summer season. Surface waters warm above 4°C around late June and cool below that temperature around mid-November, although stratification is typically unstable during early summer and fall, with large-scale hypolimnetic upwelling events occurring throughout the summer (Kozhov 1963). In the open lake, surface temperatures seldom exceed 15°C and the depth of the warm surface mixed layer is typically less than 30 m. The bays of the lake (Fig. S1) warm rapidly after ice-off in May, remain stratified for much longer than open Baikal, and reach higher summer surface temperatures (up to 20-26°C) compared to the offshore.

#### Epischura baikalensis

As the dominant pelagic consumer in Lake Baikal, *E. baikalensis* represents a key energy link in the food web of the lake (Yoshii et al. 1999; Moore et al. 2019). *E. baikalensis* completes two generations per year in Lake Baikal: a winter and a summer generation (Afanasyeva 1977). The nauplii of the winter generation hatch in late-fall through early winter and reach sexual maturity over ca. 180 days. The adults of the winter generation give rise to a summer generation, the nauplii of which hatch in July and mature over ca. 90 days, beginning to reproduce in the fall. Sexually mature *E. baikalensis* females can produce up to 10 egg sacks during their lifetime, with an average of 22 eggs per egg sack (Afanasyeva 1977). The interval between egg sacks is ca. 10 days for females of the winter generation (which reproduce in the summer) and ca. 20 days for females of the summer generation (which reproduce in late fall and winter). The majority of *E. baikalensis* individuals are found in the top 250 m layer of the water column throughout the year. During summer stratification, individuals of all ages perform diel vertical migration (DVM), with much of the population spending parts of the night in the epilimnion and descending below the thermocline during the day (Afanasyeva 1977).

#### Saprolegnia

*Saprolegnia* spp. are necrotrophic and saprotrophic fungal-like protists of the class Oomycota. Many genera of oomycetes, including *Saprolegnia*, are economically important pathogens of plants and aquatic organisms (Walker and van West 2007). *Saprolegnia* grows on and in its hosts as fine hyphae and transmission occurs through motile, biflagellate zoospores released from club-shaped sporangia (Fig. 1). Zoospores can remain viable for a few hours, undergoing repeated cycles of encystment and excystment in their search for a suitable host (Walker and van West 2007). When infecting a new host, zoospores attach externally to the host and hatch into hyphae that penetrate the integument of the host. In *E. baikalensis, Saprolegnia* infection can only be diagnosed 1–2 days after death, when hyphae break through the carapace and form a “halo” of hyphae that eventually produce sporangia and zoospores (Fig. 1).

**Figure 1:**
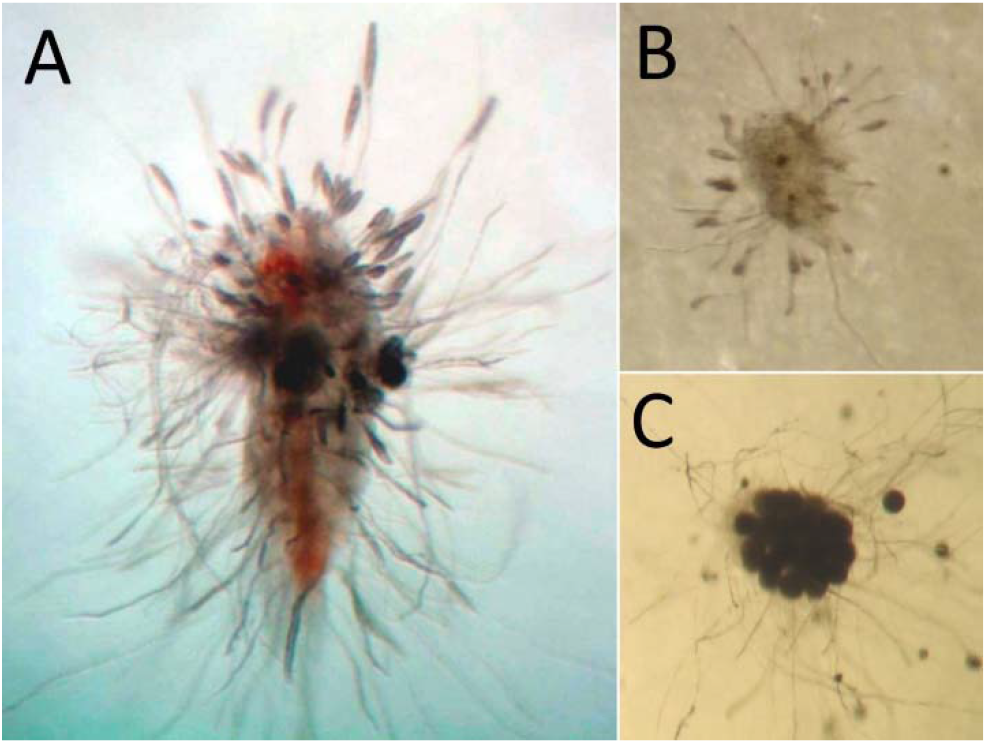
Dead *Epischura baikalensis* adult (A), nauplius (B) and egg sack (C) showing *Saprolegnia* hyphae. Club-shaped sporangia are visible in the top portion of panel A

### *Saprolegnia* isolation and culture

Growth of hyphae putatively identified as *Saprolegnia* was noted on dead *E. baikalensis* adults collected at an open water, long-term monitoring station (Stn. #1, depth ca. 800 m, Fig. S1) in the south basin of Lake Baikal in summer 2012. Infected individuals were placed on modified potato-starch, beef-broth agar at 15°C and hyphae growing from a single individual of *E. baikalensis* were isolated and grown on Sabourad agar (Wolinska et al. 2008); all subsequent work was performed with material from this strain (“Strain A”). *Saprolegnia* cultures were maintained at 15°C on a 12:12 hour light-dark cycle in incubation chambers on 35-mm petri plates. Agar plugs (5-mm diameter) from colonized plates were transferred to new agar plates every 2–3 weeks. Phylogenetic analysis of the isolated *Saprolegnia* stain (“Strain A”) was accomplished using mitochondrial-encoded Cytochrome Oxidase I (COI) and nuclear-encoded internal transcribed spacer (ITS) sequencing. Detailed methods and results for the phylogenetic analysis are provided in the supplementary information section.

### *Saprolegnia* growth on agar

Lab experiments were conducted to determine the effect of temperature on the growth of *Saprolegnia* “Strain A” on agar at 5, 10, 15, 20 and 25°C. To initiate the experiment, 5-mm circular plugs of agar and mycelia were cut from fully colonized Sabourad agar plates and placed in the center of fresh Sabourad agar plates; 3 replicate plates were used for each temperature. Plates were incubated at a 12:12 hour light-dark cycle at experimental temperatures for 72 hours; colony diameter was measured on each plate 8 times throughout the experiment to the nearest mm (as average of two perpendicular measurements). Results were expressed as average growth rate in mm/hr.

The relationship between temperature and *Saprolegnia* growth rate was explored using linear regression between average daily colony growth (mm) and temperature. Simple linear regression and 2^nd^ and 3^rd^ order polynomial regression model fits were compared using F-tests and Akaike’s Information Criterion (Crawley 2013).

### Effects of temperature and *Saprolegnia* infection on *E. baikalensis* survival and reproduction

In summer of 2013, we conducted lab experiments to assess the effect of temperature and *Saprolegnia* infection on survival and reproduction of *E. baikalensis*. Fully factorial experiments were conducted with *E. baikalensis* adult females and nauplii at 5, 10, 15 and 20°C, with and without exposure to *Saprolegnia* zoospores. *E. baikalensis* adult females and nauplii used in experiments were collected using slow zooplankton net tows from 50 m depth to the surface at an offshore station (Station #1, Fig. S1). Individuals were kept at lake surface water temperature until return to the lab, then transferred to GF/F-filtered Lake Baikal water and live-sorted to obtain enough adult females and nauplii for experiments. Only adult females with well-developed, dark ovaries and nauplii of the III-V naupliar stage were selected for the experiment. After sorting, adult females and nauplii were placed in 3-L jars with 2.5 L of GF/F-filtered lake water at ambient surface water temperature and the jars were placed in temperature-controlled chambers at 5, 10, 15 and 20°C for 4 hours to slowly bring animals to experimental temperatures.

After the 4-hour acclimation, live adults were transferred into 6-well plates with 6 mL of GF/F-filtered water at experimental temperature in each well (1 individual per well); nauplii were transferred to 24-well plates with 2 mL of GF/F filtered water per well. Plates were divided into 3 groups (18 adults or 24 nauplii per group): a *Saprolegnia* treatment group and two control groups – a *Saprolegnia* sporulation medium control group and a “true” control group. Wells in the *Saprolegnia* treatment group received 0.4 mL/well of *Saprolegnia* sporulation medium (see below) with active zoospores, aiming for a final concentration of 60 zoospores/mL. Wells in the sporulation medium control group received 0.4 mL/well of filtration-sterilized (triple filtration through 0.2-μm syringe filters) sporulation medium and wells in the “true” control group received 0.4 mL of GF/F-filtered lake water. The sporulation medium control group was used to account for potential negative effects of the sporulation medium or dissolved toxins produced by *Saprolegnia* on *E. baikalensis* survival and reproduction. Zoospore production by cultured *Saprolegnia* was induced using methods similar to those described in Wolinska et al. (2008). Briefly, 5-mm plugs of agar from fully-colonized Sabourad agar plates were placed in 100 mL of GF/F-filtered and boil-sterilized water at 15°C. The sporulation medium was inspected under a compound microscope for presence of zoospores and replaced every 24 hours until sporangia and live (actively swimming) zoospores were observed. Spore concentrations were determined by counting subsamples of the sporulation medium preserved and stained with Lugol’s iodine in a hemocytometer.

All experiments were checked at least every 24 hours (more frequently during the first few days of the experiment). The following information was recorded about all adult females at each check: live or dead, presence of visible *Saprolegnia* growth, presence of egg sack, presence of *Saprolegnia* hyphae on egg sack, and presence and number of newly hatched nauplii. In experiments with nauplii, we recorded whether individuals were live or dead and whether *Saprolegnia* hyphae were present. To minimize handling of *Epischura* and to reduce the possibility of algal overgrowth or algal medium toxicity to *E. baikalensis* during the experiment, animals were not fed and water was not replaced during experiments.

We used parametric survival analysis to assess the effect of *Saprolegnia* infection status and temperature on the survival *of E. baikalensis* adults and juveniles. Prior to analysis, we determined whether survival differed significantly between “true” control treatments and *Saprolegnia* sporulation medium control treatments. No differences in survival between the two controls were found for either adults or juveniles, so we used the data from both controls treatments as the control group for subsequent analyses. Choice of error distribution for survival analysis was assessed using Akaike Information Criterion (AIC) and model comparison (Crawley 2013; George et al. 2014); exponential or Weibull distributions were identified as most appropriate in all cases (see results for details). Survival models were parameterized with temperature, infection status and their interaction as predictor variables. Survival analysis was implemented using the *survreg()* function in the ‘survival’ package for R (version 2.40-1; Therneau and Lumley 2014). Predicted survival times for use in the population model (see below) were extracted using the *predict()* function.

Effects of temperature and *Saprolegnia* exposure on *E. baikalensis* reproductive parameters were tested using ANOVA (Type III sum of squares) with temperature, infection status and their interaction as predictor variables. The response variables were time from start of the experiment to production of 1^st^ egg sack, time from egg sack production to hatching of 1^st^ nauplii, and total number of live nauplii produced per egg sack. Prior to conducting the analysis, we tested whether any of the three response parameters differed significantly between the two control treatments. We found no differences for any of the response variables, so data from both control treatments were used as the control group in subsequent analyses. In the *Saprolegnia* exposure treatment, we observed that some egg sacks developed visible hyphae and some did not. Consequently, we carried out the statistical analysis twice: once without separating egg sacks in the *Saprolegnia* treatment based on whether they showed hyphae and once separating them based on this parameter.

### Long term data

To examine the seasonal dynamics of *Saprolegnia* infection prevalence in *E. baikalensis* as well as the seasonal dynamics of *E. baikalensis* abundance in relation to temperature, we used long term monitoring data. Data were collected by researchers from Irkutsk State University at a pelagic station (Station #1; Fig. S1) in the southern basin of Lake Baikal between 1945 and 2003 at a sub-monthly frequency. Samples were collected with zooplankton closing nets; typical depth intervals for sampling were 0–10, 10–25, 25–50, 50–100, 100–150, 150–250 and 250–500 m. These rich data have recently been used to examine long-term trends in the ecology of the lake (Hampton et al. 2008; Izmest’eva et al. 2011; Izmest’eva et al. 2016; Silow et al. 2016) and additional information about sampling and data is available in those papers.

Data on presence of visibly Saprolegnia-infected *E. baikalensis* were recorded for 16 years. It is unknown whether *Saprolegnia*-infected individuals were recorded on every occasion they were present (i.e., *Saprolegnia* infection was only present in 16 years between 1945 and 2003), or at the discretion of researchers working with the samples in particular years. However, it appears that in years when *Saprolegnia* was noted, the number of individuals with infection was counted in all samples such that the data robustly indicate the prevalence of infection at each depth and year. Given the uncertainty associated with sample processing, these data should be interpreted with caution, but still enable some insight into seasonal and depth patterns of infection prevalence. We used these data to examine patterns in the depth distribution of infection prevalence for the 0–500 m layer, seasonal patterns in infection prevalence, and the relationship between surface water temperature and infection prevalence. It is important to point out that infection can only be diagnosed reliably after death, once *Saprolegnia* hyphae break through the body wall and become visible. Additionally, there is a delay (1–2 days in our experiments) between death and appearance of visible hyphae. Thus, these data provide somewhat indirect and probably conservative estimates of infection prevalence.

To assess seasonal patterns in the abundance of *E. baikalensis* in relation to surface temperature, we analyzed 52 years of the long-term data using generalized additive mixed models (GAMMs). We separated *E. baikalensis* individuals into juveniles (nauplii and copepodites) and adults, expressing densities as number of individuals per m^2^ in the 0–250 m layer. Maximum monthly summer temperatures in the 0–25 m layer were used to classify years as ‘cold’ (0–10^th^ percentile), ‘cool’ (10^th^–35^th^ percentile), ‘average ‘(35^th^–65^th^ percentile), ‘warm’ (65^th^–90^th^ percentile) and ‘hot ‘. GAMMs of abundance vs. day of year (DOY) were used to fit curves for years in each temperature category, with individual years as a random factor using the ‘mgcv’ package in R (version 1.8-12; Wood 2001). We created separate models for adults and juveniles and, for each age category, separate models for full year abundance (DOY 1–365) and for the period where surface temperatures typically exceed 4°C (DOY 180–320 across the time series), which we defined as the “stratified season”. We asked whether there was a significant seasonal variation in densities and whether the pattern of variation differed with year types (‘cold’, ‘cool’, etc.). *E. baikalensis* densities were cube root transformed to satisfy assumptions of normal distribution of residuals and equal variance.

We also examined the effect of summer temperatures on *E. baikalensis* adult and juvenile densities using linear regression analysis. We regressed average “stratified season” (DOY 180–320) densities of adults and juveniles (cube root-transformed) against maximum annual surface water (0–25 m layer) temperatures. We also regressed densities of adults and juveniles in fall and early winter (DOY 320–365) against maximum annual temperatures to assess how “stratified season” temperature affected density in the following season.

### Model

To examine the population-level effects of water temperature and *Saprolegnia* infection on *E. baikalensis*, we constructed a population model for *E. baikalensis* (see supplementary information section for detailed explanation of model; Fig. S2). The model included healthy (uninfected) adults, infected adults, healthy juveniles and infected juveniles. The model was parameterized using a combination of experimentally-determined and literature-estimated values for life stage-and infection status-specific mortality (m), reproduction (r), maturation (g), and infection transmission rates (β) (Table 1, supplementary information section). We used long-term surface water (0–25 m) temperature data (1948–2002) from a pelagic station in Lake Baikal to construct seasonal temperature scenarios representative of ‘cold’, ‘cool’, ‘average’, ‘warm’ and ‘hot’ years in the pelagic zone (same definitions of temperature categories as in the long-term data analysis). We also used seasonal temperature data to simulate conditions in a warm shallow bay (Proval Bay) and a large, deep bay (e.g., Barguzin or Chivyrkuy Bay). To examine the effect of DVM on *E. baikalensis* populations under different temperature and infection scenarios, we created models where *E. baikalensis* spent half of the day in the hypolimnion and models where *E. baikalensis* were restricted to the epilimnion (as might be the case in shallow regions of the lake).

**Table 1:**
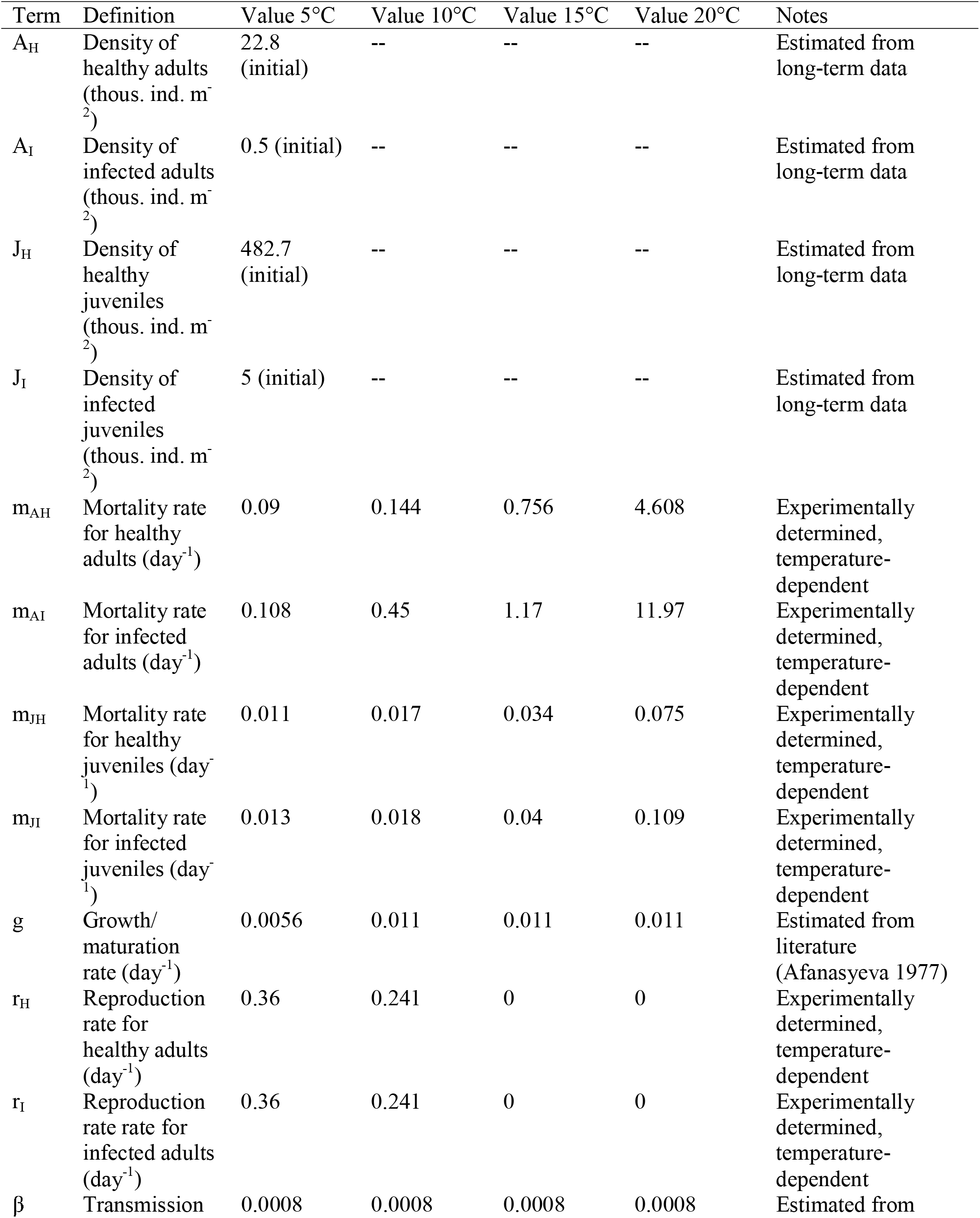
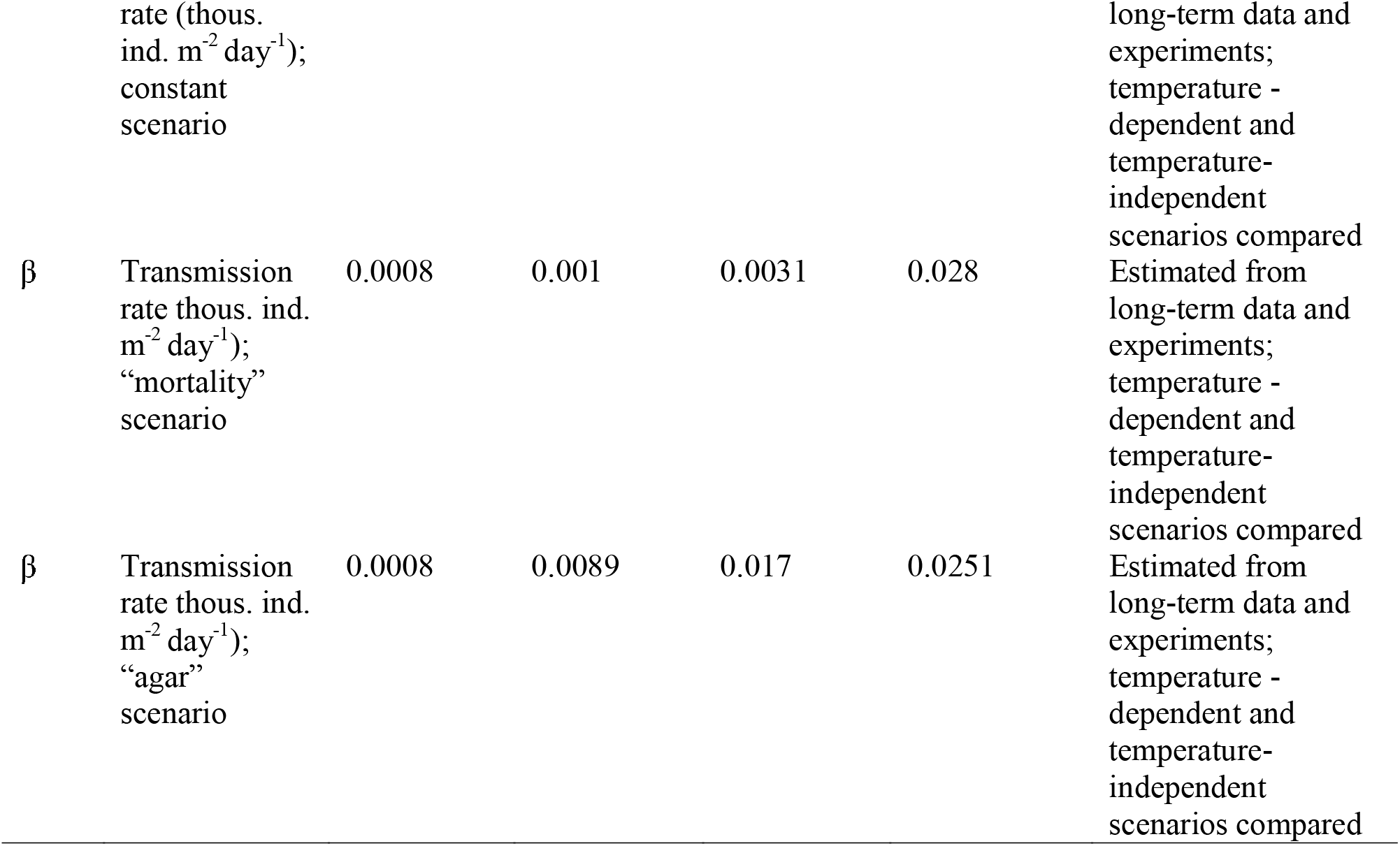
Terms and values used in population model. For details on infection rate scenarios, see methods and supplementary information sections.

The infection rate parameter (β) is difficult to determine even for well-studied systems, and the choice of β can have large effects on model outcomes (Kirkeby et al. 2017). Given that the *E. baikalensis – Saprolegnia* host-parasite relationship has received no study, the choice of the value for β used in the model was difficult. We therefore compared the output of model scenarios parameterized with three different β-temperature functional relationships. The first scenario assumed a temperature-independent β(“temperature-invariant β” scenario). In the other two β scenarios increased with temperature; in one the β-temperature relationship was based on the temperature-specific growth rates of *Saprolegnia* on agar (“agar β” scenario) and in the second on the change in mortality rates of *Saprolegnia-infected E. baikalensis* at different temperatures (“mortality β” scenario). Additional details on these scenarios is presented in the supplementary information section.

Since our primary interest was to explore the effect of temperature variation on *E. baikalensis* populations, models were ran only for the duration of the stratified summer season, when surface temperatures were >4°C. This period differed for different temperature scenarios, ranging from DOY 170–330 (Jun. 19–Nov. 27) in the “cold pelagic” scenario to DOY 129-303 (May 9–Nov. 1) in the “Proval Bay” (warm water) scenario.

## Results

### *Saprolegnia* identification and growth rate

COI-and ITS-based phylogenies confirm that the oomycete parasite infecting *E. baikalensis* belongs to the genus *Saprolegnia* (Fig. S3). Mismatches between phylogenetic relationships and species boundaries (i.e., for several *Saprolegnia* species, strains of the same species did not form monophyletic groups) make it difficult to unambiguously identify our strain to species level. However, both the single-gene ITS and COI trees and the phylogeny reconstructed from concatenated data (either with IQtree or RaxML) suggest the parasite is closely related to *S. diclina* (see supplementary information for more detail).

*Saprolegnia* growth rate increased significantly with temperature (simple linear model, R^2^=0.93, p<0.001). A simple linear fit was more parsimonious than 2^nd^ or 3^rd^ order polynomials for the temperature - *Saprolegnia* growth rate relationship, although visual inspection of results suggests that growth rates may have begun reaching a plateau near 25°C (Fig. 2).

**Figure 2:**
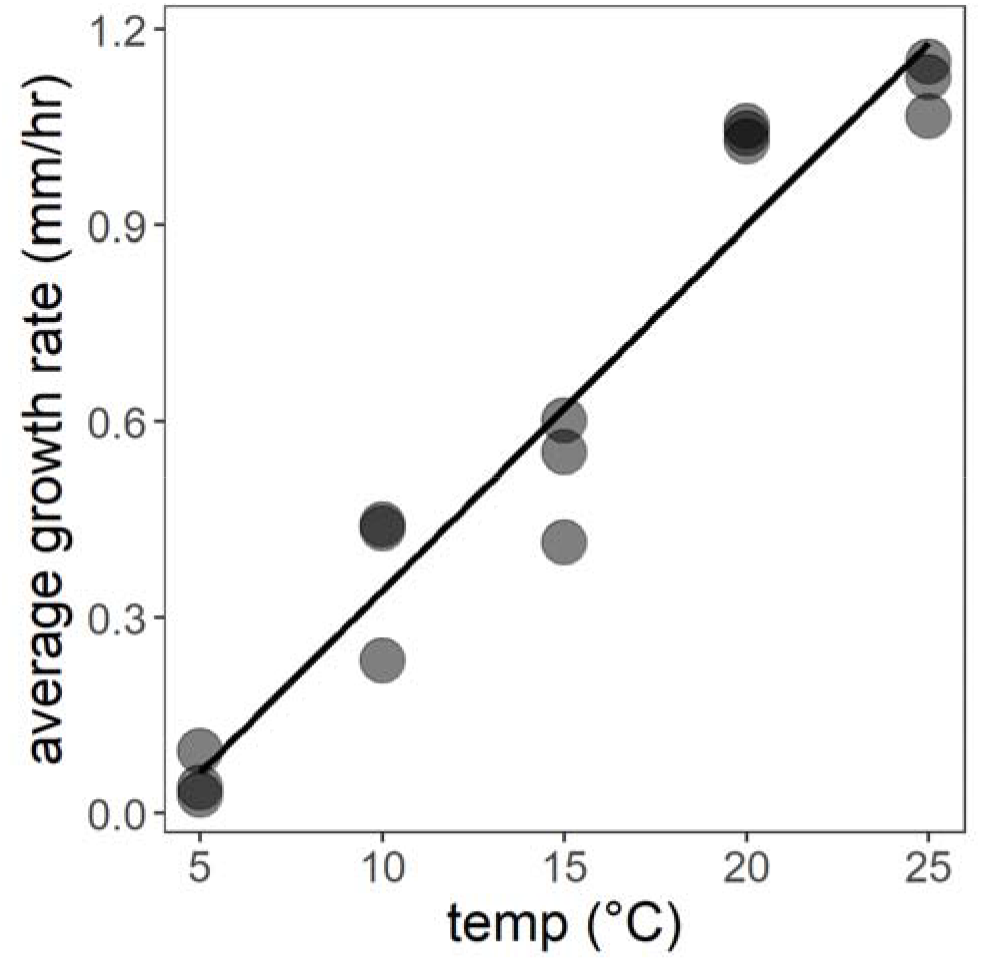
Growth rates of *Saprolegnia* on Sabourad agar at experimentally manipulated temperatures (5, 10, 15, 20, and 25°C), with linear regression fit shown (R^2^ = 0.93).

### Survival of *E. baikalensis*

Temperature and *Saprolegnia* infection status significantly affected survival rate of *E. baikalensis* nauplii and adults, with no statistically significant interaction between the two terms (Table 2). Survival decreased with increasing temperature and exposure to *Saprolegnia* (Fig. 3; Table S1). None of the individuals that died in control treatments showed growth of *Saprolegnia* hyphae after death. In contrast, many individuals that died in the *Saprolegnia* treatment showed visible growth of *Saprolegnia* hyphae after death (40% across all temperatures), with the proportion of individuals showing growth of hyphae increasing with temperature (Table 3). For adult and juvenile individuals that showed *Saprolegnia* growth after death, the average time between death and visible development of external hyphae averaged 1.26 (±1.26 SD) days across all temperatures (Table 3).

**Figure 3:**
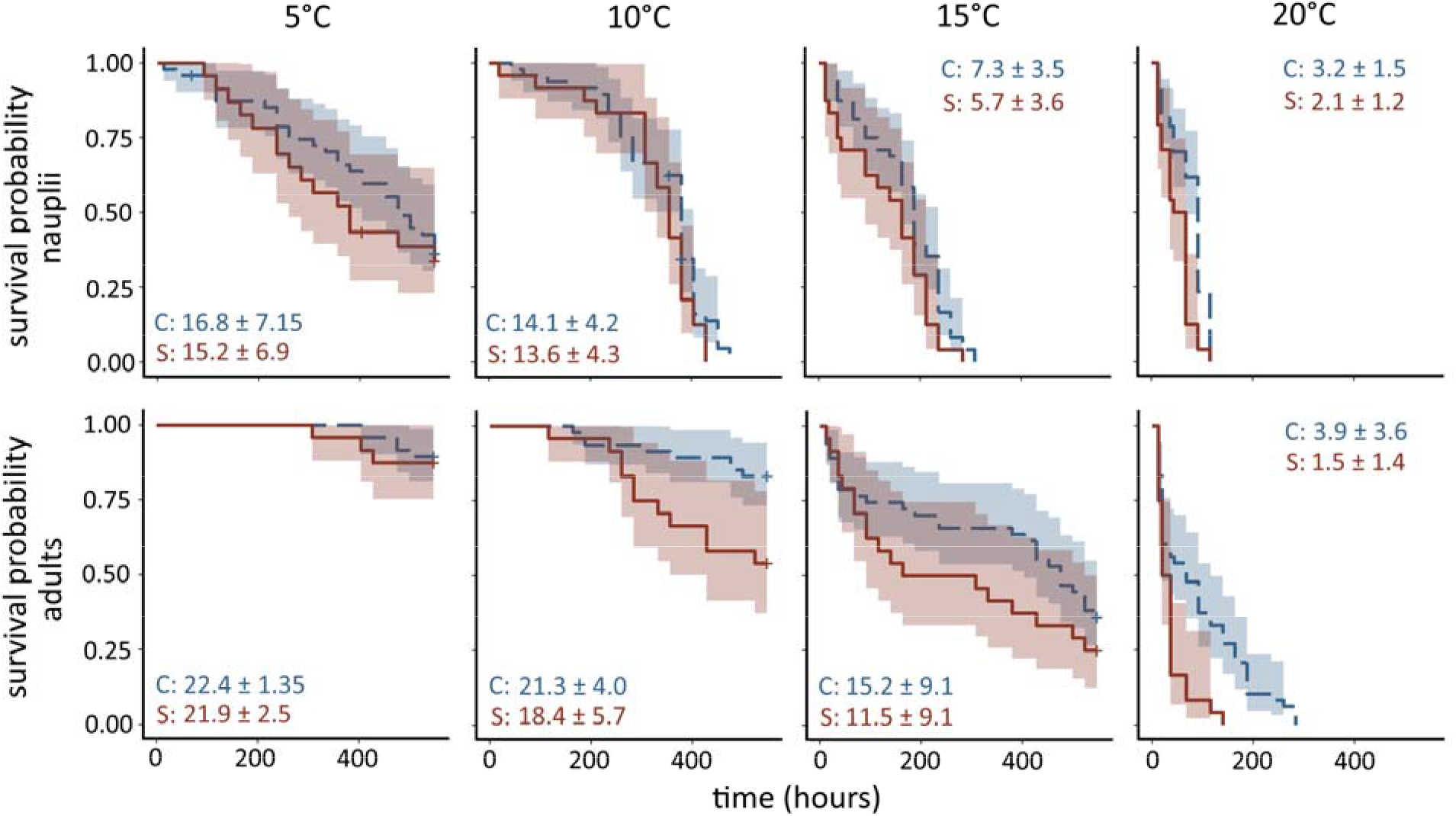
Survival curves of *Epischura baikalensis* nauplii (top panels) and adults (lower panels panels) in control (blue dashed lines) and *Saprolegnia* exposure treatments (red solid lines) at 5, 10, 15 and 20°C. + symbols indicate censoring events (loss of individual from experiment not due to death or survival to end of experiment). Shaded regions represent 95 % confidence intervals. Numbers represent average survival times in days (± SD) for control (C) and *Saprolegnia*-exposed (S) groups.

**Table 2:** Results of parametric survival analysis for *Epischura baikalensis* nauplii and adults, examining effect of temperature and *Saprolegnia* exposure treatments.

| Group | Probability distribution | p-value |
| --- | --- | --- |
| *Epischura* nauplii | Weibull |  |
| Temperature |  | <0.001 |
| Treatment |  | 0.011 |
| Interaction |  | 0.12 |
| *Epischura* adults | Exponential |  |
| Temperature |  | <0.001 |
| Treatment |  | <0.001 |
| Interaction |  | 0.71 |

**Table 3:**
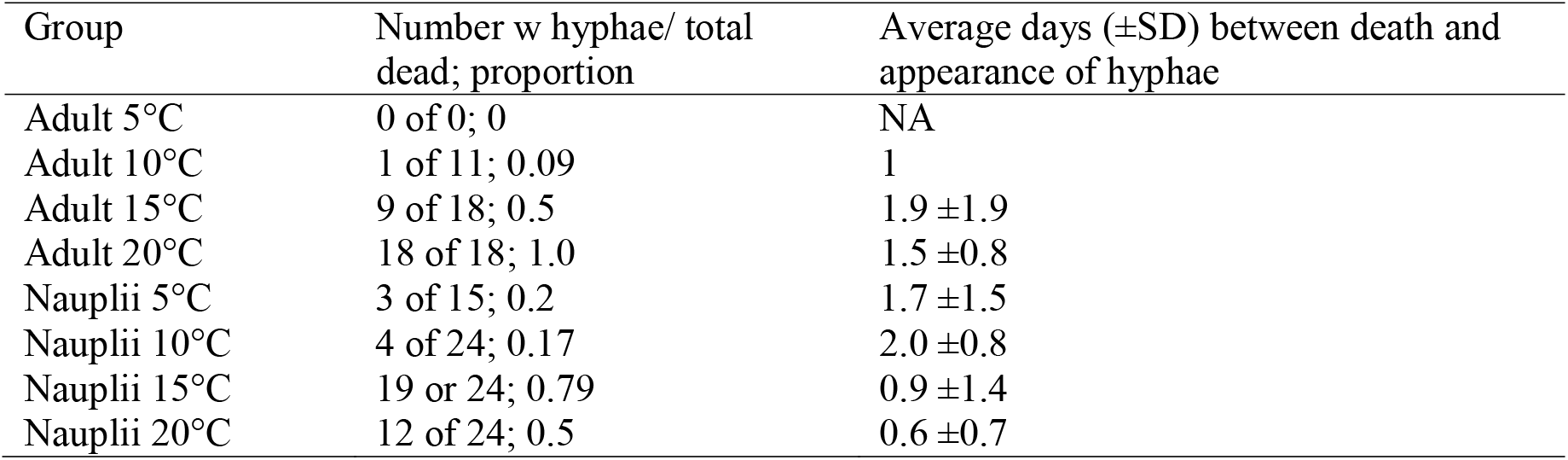
Proportion of *Epischura baikalensis* adults and nauplii that showed *Saprolegnia* growth after death and average time between death and noticeable *Saprolegnia* growth.

### Reproduction of *E. baikalensis*

Temperature and *Saprolegnia* exposure had different effects on reproductive parameters of *E. baikalensis* (Fig. 4; Table 4; Table S2). Time to production of first egg sack was not affected by *Saprolegnia* exposure and decreased significantly with increasing temperature; however, warming temperature ultimately limited egg production, as no egg sacks were produced at 20°C. Time between egg sack production and hatching was also unaffected by *Saprolegnia* exposure and decreased significantly from 5°C to 10°C; no eggs produced at 15°C and 20°C hatched. Number of nauplii hatching from egg sacks varied both with temperature and *Saprolegnia* exposure. Results of analysis differed depending on whether we combined results for all *Saprolegnia* exposed adults or separated them based on presence of hyphae on egg sacks. When combining results of all *Saprolegnia-exposed* adults we saw a negative effect of increasing temperature on number of hatched nauplii, a significant positive effect of *Saprolegnia* exposure on number of hatched nauplii and a significant interaction between the predictors, where the positive effect of *Saprolegnia* exposure was larger at 5°C than 10°C. When Saprolegnia-exposed individuals were separated based on presence of hyphae on egg sacks, increasing temperature still had a significant negative effect on number of live nauplii, but there were significant differences depending on whether hyphae were present or not on egg sacks of *Saprolegnia*-exposed adults. Egg sacks of individuals exposed to *Saprolegnia*, but not showing hyphal growth, yielded the most nauplii, followed by egg sacks showing hyphal growth. Egg sacks of control individuals yielded the fewest nauplii. This pattern was not strongly different across temperature (interaction p=0.06).

**Figure 4:**
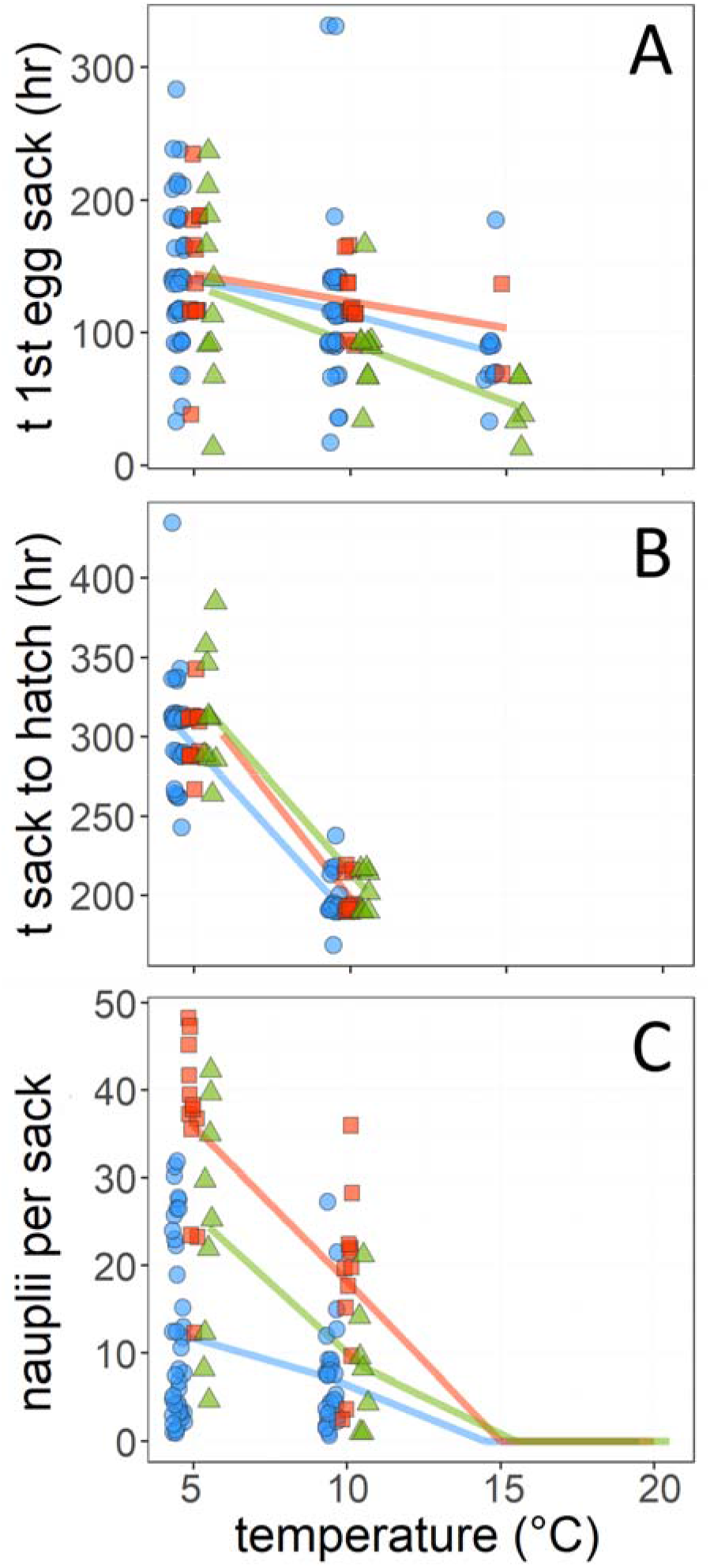
Effect of temperature and *Saprolegnia* on reproduction of *Epischura baikalensis*, including time to production of 1^st^ egg sack (A), time from egg sack production to first hatch (B) and number of live nauplii per egg sack (C). Blue circles= control treatments, red squares= individuals exposed to *Saprolegnia* spores but not showing development of hyphae on egg sack, green triangles= individuals exposed to *Saprolegnia* spores and showing development of hyphae on egg sack. Lines connect average values within categories.

**Table 4:** Results of ANOVA on effect of temperature and *Saprolegnia* exposure on *Epischura baikalensis* reproductive parameters. Results for two analyses are shown: combining all results from Saprolegnia-exposed adults (Treatment: control vs. Saprolegnia-exposed) and separating *Saprolegnia-exposed* adults based on whether visible hyphae were present on the egg sack (Treatment: control vs. Saprolegnia-exposed without hyphae vs. Saprolegnia-exposed with hyphae). Tukey HSD codes: 5, 10, 15: temperature (°C) treatments, C: control, S: all *Saprolegnia-exposed* adults, SN: *Saprolegnia-exposed*, but no hyphae on egg sack, SH: *Saprolegnia-exposed* and hyphae present. Treatments with different superscripted letters are significantly different (p < 0.05) according to the Tukey HSD test.

|  | Combined *Saprolegnia* results |  |  |  | Separated by hyphal presence |  |  |  |
| --- | --- | --- | --- | --- | --- | --- | --- | --- |
|  | df | F-value | p-value | Tukey HSD | df | F-value | p-value | Tukey HSD |
| (Time to 1<sup>st</sup> egg sack)<sup>0.5</sup> |  |  |  |  |  |  |  |  |
| Temperature | 2 | 7.36 | 0.0009 | 5<sup>a</sup>, 10<sup>b</sup>, 15<sup>c</sup> | 2 | 6.32 | 0.0023 | 5<sup>a</sup>, 10<sup>b</sup>, 15<sup>c</sup> |
| Treatment | 1 | 0.22 | 0.64 |  | 2 | 0.98 | 0.38 |  |
| Interaction | 2 | 0.77 | 0.47 |  | 4 | 1.18 | 0.32 |  |
| Time to hatch |  |  |  |  |  |  |  |  |
| Temperature | 1 | 146.18 | <<0.0001 | 5<sup>a</sup>, 10<sup>b</sup> | 1 | 146.69 | <<0.0001 | 5<sup>a</sup>, 10<sup>b</sup> |
| Treatment | 1 | 0.029 | 0.96 |  | 2 | 0.12 | 0.89 |  |
| Interaction | 1 | 0.01 | 0.92 |  | 2 | 0.35 | 0.71 |  |
| (Number of nauplii)<sup>0.5</sup> |  |  |  |  |  |  |  |  |
| Temperature | 1 | 4.1 | 0.046 | 5<sup>a</sup>, 10<sup>b</sup> | 1 | 4.40 | 0.038 | 5<sup>a</sup>, 10<sup>b</sup> |
| Treatment | 1 | 39.16 | <<0.001 | C<sup>a</sup>, S<sup>b</sup> | 2 | 23.29 | <<0.0001 | C<sup>a</sup>, SN<sup>b</sup>, SH<sup>c</sup> |
| Interaction | 1 | 4.9 | 0.029 |  | 2 | 2.92 | 0.058 |  |

### *Saprolegnia* prevalence in *E. baikalensis* populations

Results from years in which *Saprolegnia* presence was recorded in the long-term data reveal several patterns related to *Saprolegnia* infection prevalence (Fig. 5). First, while the majority of *E. baikalensis* individuals are concentrated in the top 100 m of the water column, the abundance of visibly *Saprolegnia-infected* individuals peaks lower in the water column (100–150 m), and the prevalence of infected individuals increases with depth (Fig. 5A–C). Second, infection prevalence seems to increase with temperature, peaking in late summer and early fall (Fig. 5D–E). More than 5% of the population in the 0–500 m layer were visibly infected on several dates between August and November.

**Figure 5:**
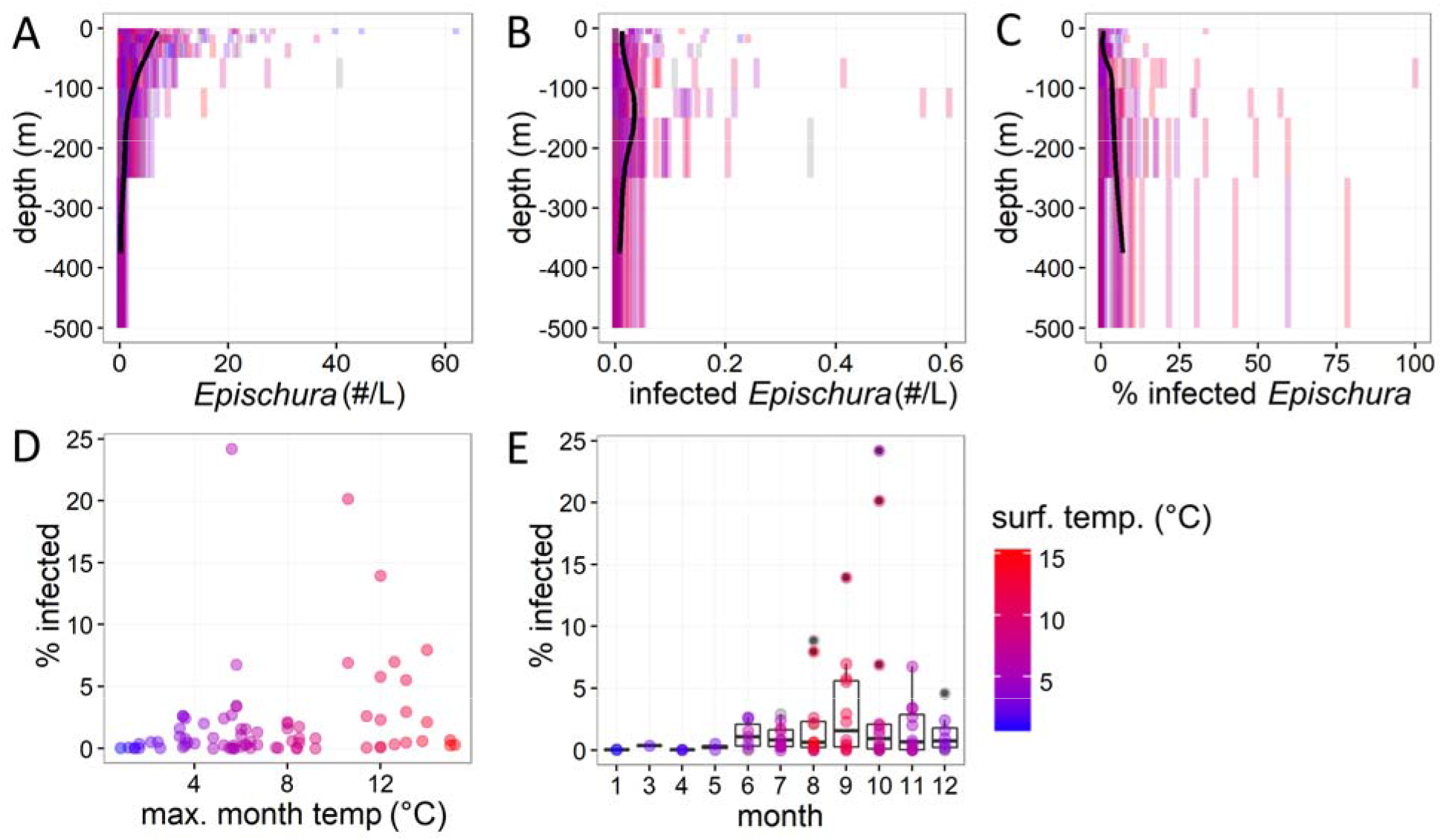
Patterns of *Saprolegnia* infection in pelagic zone of Lake Baikal based on 16 years of data (collected 1945–1999) in which *Saprolegnia* presence was recorded. Depth distribution of A) total *Epischura* abundance, B) number of visibly infected *Epischura* per liter, and C) prevalence of visibly infected *Epischura* plotted across all sampling dates. Black lines in A-C represent a LOESS smoother. D) Prevalence of visibly infected *Epischura* in the 0–500 m layer plotted against max. monthly temperature in the 0–50m water layer. E) Prevalence of visibly infected *Epischura* in the 0–500 m layer by month. In panels A, B and C vertical bars represent densities for net hauls from different depth intervals. Color corresponds to maximum monthly temperature in top 50 m for each sample.

### Seasonal patterns of *E. baikalensis* abundance

GAMM analysis showed that abundance of adult *E. baikalensis* varied significantly through the year as well as through the stratified season, but without a significant difference in the seasonal pattern of fluctuation between years in different temperature categories (Table S3). Across all years, *E. baikalensis* adults increased in abundance through winter and spring, decreasing in abundance after the onset of stratification (Figs. 6, S4). Juvenile *E. baikalensis* (nauplii and copepodites) also varied in abundance though both the entire year and within just the stratified season, increasing through winter and peaking in summer, later than adult densities (in August). There was a significant difference in the pattern of variation in juvenile densities in years in different temperature categories, with apparently earlier and larger peaks in abundance in warmer than cooler years (Table S3; Figs. 6, S4).

**Figure 6:**
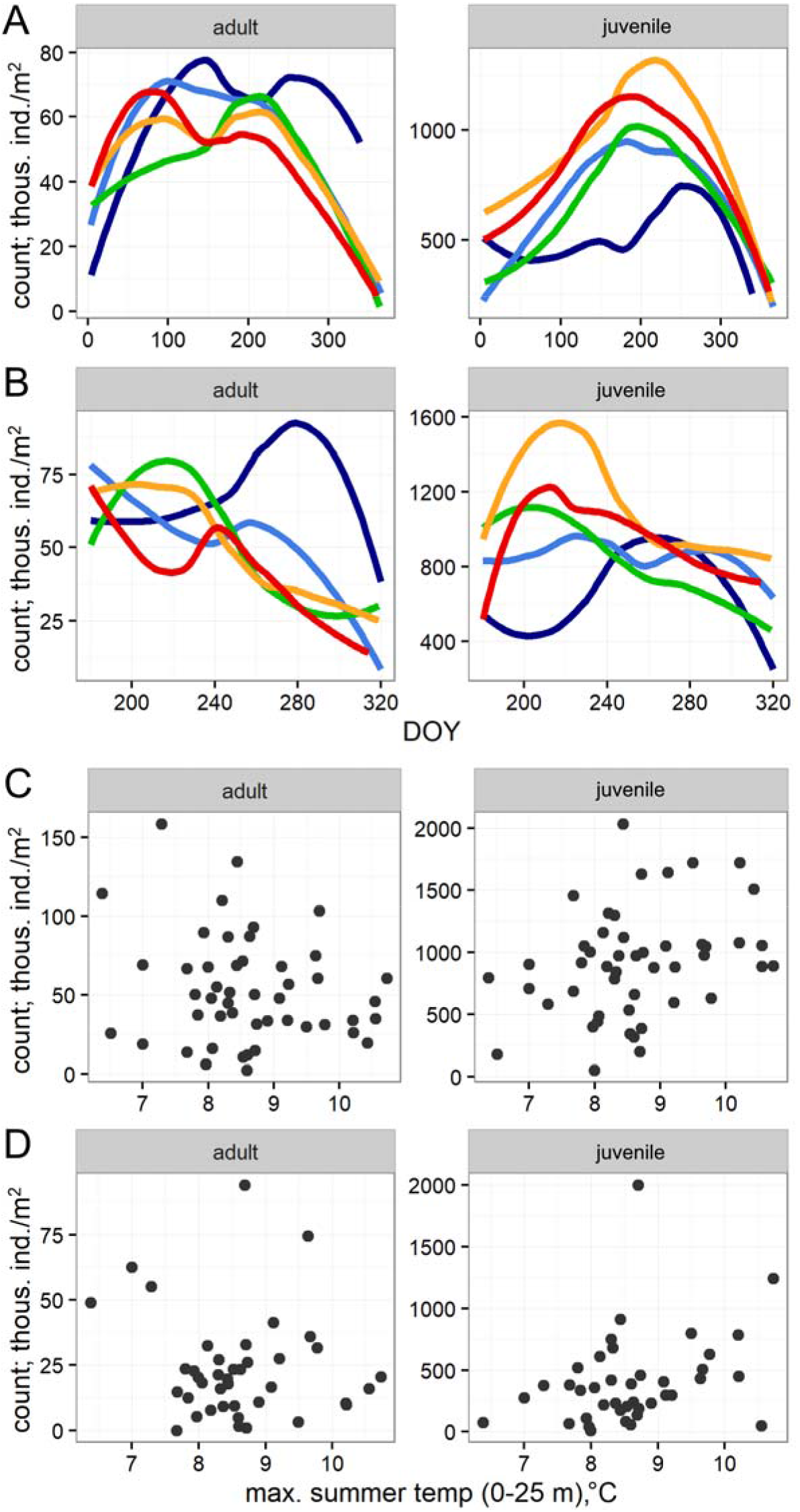
Seasonality and temperature dependence of adult and juvenile *Epischura* abundance in 0–250 m layer in pelagic zone of Lake Baikal based on 48 years of data. A) LOESS smoothers fitted to year-round densities in cold (dark blue), cool (light blue), average (green), warm (orange) and hot (red) years (see results section for detail on year classification). B) same as in A but for stratified season only. C) annual average summer-time (DOY 180–320) densities of *Epischura* plotted against max summer lake surface temperatures. D) annual average fall-winter (DOY > 320) densities of *Epischura* plotted against max summer surface temperatures.

Regression analysis showed no significant relationship between maximum annual lake surface temperature and the stratified season abundance of adults (R^2^=0.0, p=0.38) and a weak but significant positive relationship between lake surface temperature and juvenile densities during the stratified season (R^2^=0.11, p=0.02). A similar pattern (Fig. 6) occured for the relationship between maximum temperature and fall/early winter densities of adults (R^2^=0.11, p=0.55) and juveniles (R^2^=0.09, p=0.03).

### Model

Our population model generally reproduced the seasonal pattern of *E. baikalensis* abundance and correctly predicted the timing of peak *Saprolegnia* prevalence during late summer (Figs. 5–7). The model also correctly predicted the absence of *E. baikalensis* from shallow bays, a prediction that is supported by observations reported in the literature (e.g., Kozhov 1963, Bowman 2014). On the other hand, the model overestimated the negative effect of warm pelagic temperatures on *E. baikalensis* densities; while the model predicted large decreases in *E. baikalensis* abundance from cold to hot years, long-term observations show a slight positive relationship between summer surface temperatures and late fall *E. baikalensis* densities in the pelagic region of the lake (Fig. 6).

**Figure 7:**
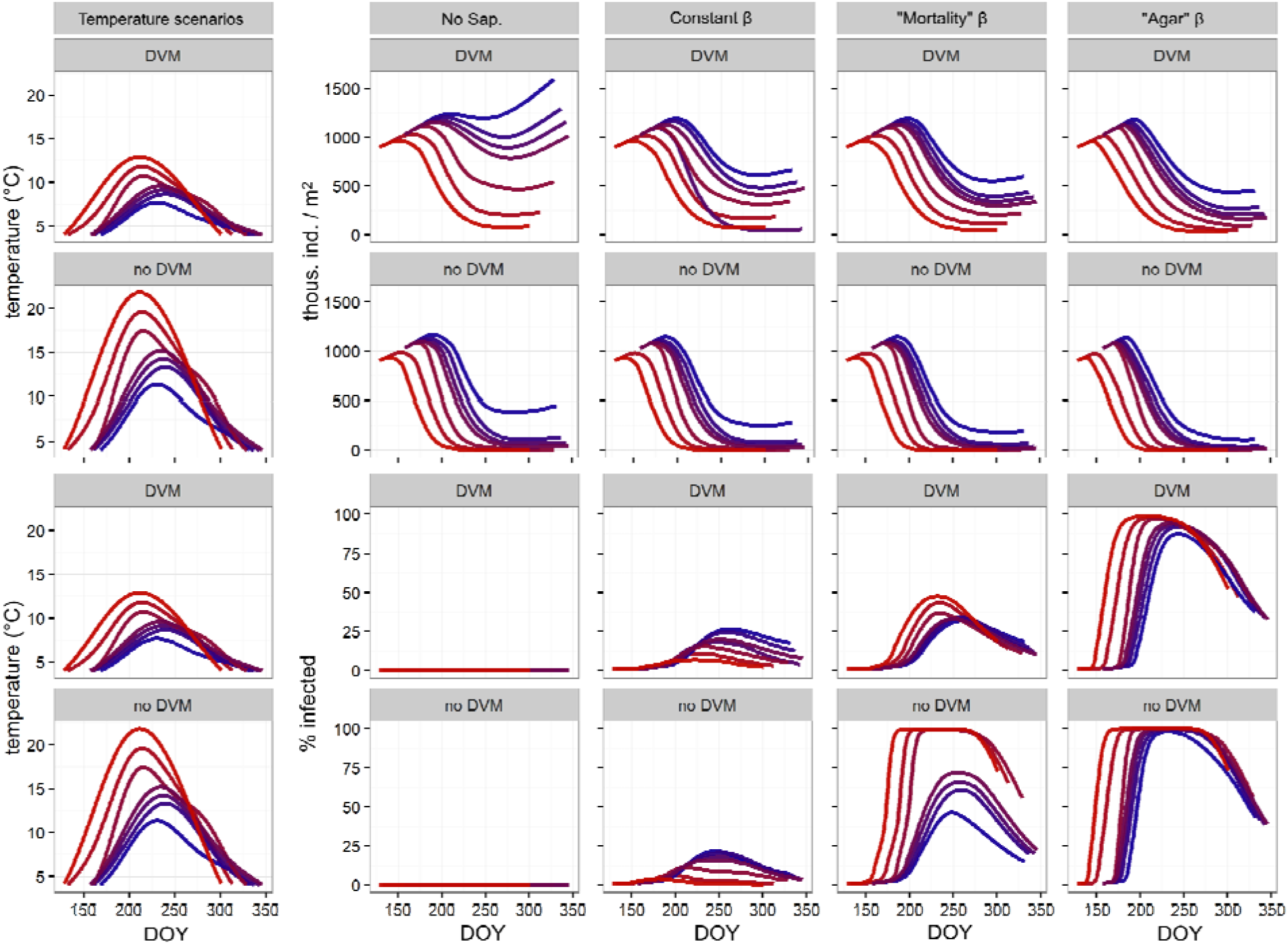
Summary of model output showing the effect of water temperature and *Saprolegnia* infection on total abundance of *Epischura* (adults and juveniles) and percent of individuals infected with *Saprolegnia* for the stratified period under different temperature scenarios. Temperature scenarios (from coldest to hottest) include cold, cool, average, warm and hot years in the pelagic zone, and typical years in a large bay and a small shallow bay of Lake Baikal. Different colors correspond to different temperature scenarios (blue= coldest, red= warmest). Model runs were performed for theoretical populations performing DVM and theoretical populations not performing DVM. Four different *Saprolegnia* infection scenarios were examined: one with no *Saprolegnia* present, and three with different infection rate (β) – temperature relationships. Details of the β – temperature relationships used in the model can be found in the text.

Across all modeled temperature scenarios (cold to hot years in pelagic, and typical years in deep and shallow bays), our temperature-driven population model predicted an S-shaped seasonal pattern of *E. baikalensis* abundance during the stratified season (Figs. 7, S5–S8). Abundances increased during the first part of the stratified season, followed by a decrease during peak temperatures and, in the cooler scenarios, an increase in abundance during the final part of the stratified season as surface temperatures decreased. The model predicted large differences in *E. baikalensis* abundance among the different temperature and infection scenarios, with a strong pattern of decreased *E. baikalensis* abundance with increasing surface temperatures. Scenarios where *E. baikalensis* were able to perform DVM had higher predicted *E. baikalensis* abundances across all temperature and *Saprolegnia* scenarios. Several model scenarios predicted extinction of *E. baikalensis* during the open-water period; extinction was predicted for the warmer scenarios, where *E. baikalensis* were unable to perform DVM (Fig. 7). DVM allowed *E. baikalensis* to persist (albeit at low densities) even under the warmest scenarios.

The presence of *Saprolegnia* had a negative effect on *E. baikalensis* densities across all temperature scenarios, although the magnitude of the effect varied with the relationship between the infection rate parameter (β) and temperature. The shape of the relationship between β and temperature also had a large effect on modeled infection prevalence. The “temperature-invariant β” scenario predicted the lowest overall infection prevalence of the three scenarios and, unlike the other two scenarios, predicted higher rates of infection prevalence for the lower temperature scenarios. On the other hand, the two scenarios where β increased with temperature (“mortality” and “agar” scenarios) predicted higher overall infection prevalence, as well as a positive relationship between infection prevalence and temperature. The non-linear relationship between β and temperature in the “mortality” scenario resulted in interesting differences for predicted infection prevalence between different temperature scenarios. Under the “mortality” and “agar” β scenarios, infection prevalence could be high, with several scenarios predicting infection of the majority of the population. Interestingly, the scenarios with high infection rate prevalence did not result in dramatically different predictions of *E. baikalensis* abundance compared to similar temperature scenarios with lower infection rates.

## Discussion

This study provides a detailed description of a previously unstudied host-parasite system and illustrates how multiple stressors affect a cold-adapted dominant organism in one of the world’s largest lakes. Results of experiments, modeling and long-term monitoring suggest that the thermal mismatch (Cohen et al. 2017) between *E. baikalensis* hosts and their oomycete parasites can lead to negative synergistic effects on the hosts under warming temperatures. Modeling also shows that DVM can provide an important thermal refuge to *E. baikalensis* from both the direct negative effects of temperature and the impacts of parasitism. These results are broadly relevant to understanding how future climate warming will affect pelagic ecosystems of cold lakes and oceans.

### Effects of temperature and parasitism: experimental results

Our experimental studies show a large mismatch between the thermal optima of *E. baikalensis* and their oomycete parasite *Saprolegnia* (Figs. 2–4). These results support our hypotheses (H1-H3) that increasing temperatures will decrease fitness of *E. baikalensis*, both directly and via the negative effects of *Saprolegnia*. Studies on various strains of *Saprolegnia* have consistently shown that the thermal optimum for this genus occurs around 25°C (Oláh and Farkas 1978, Hatai et al. 1990). Our results closely match these findings, showing highest growth rates on agar at 25°C. The thermal optimum for *E. baikalensis* is nearer to 5°C; survival was highest at this temperature for both adults and juveniles, and average survival times decreased rapidly at 15 and 20°C compared to 5 and 10°C. Temperature also had a dramatic effect on the fitness of *E. baikalensis* through effects on reproductive output, with complete reproductive failure at 15 and 20°C. While we are not aware of other published experimental results for *E. baikalensis*, Kozhov (1963) and Afanasyeva (1977), both cite field observations showing that *E. baikalensis* does not tolerate temperatures above 15°C.

An unexpected finding that contradicts our hypotheses about the impact of *Saprolegnia* (H3) was the apparent increase in reproductive output (as number of nauplii hatching per egg sack) of *E. baikalensis* exposed to *Saprolegnia* (Fig. 4). We are unsure whether this effect is a function of increased number of eggs per egg sack in Saprolegnia-exposed individuals or of lower egg mortality rates of exposed females. This result is especially surprising given that the few other studies that examined effects of oomycete parasites *(Aphanomyces* spp.) on freshwater copepods have found high brood mortality associated with parasitism (Burns 1989; Miao and Nauwerck 1999; Valois and Burns 2016). This finding also contrasts with most other studies on the effect of parasites in terrestrial and aquatic host-parasite systems, where negative impacts of infection on fecundity are typical (Stirnadel and Ebert 1997; Hurd 2001, Ebert 2005; Duffy and Hall 2008; Cohen et al. 2017).

One possible explanation for the increased reproductive output of infected *E. baikalensis* individuals is compensation by infected mothers for their infection-associated shorter lifespan, an idea known as the “terminal investment hypothesis” or “fecundity compensation”. Results congruent with fecundity compensation have been demonstrated for diverse organisms infected with parasites or diseases, including snails, birds, amphibians and freshwater *Daphnia* (Minchella and Loverde 1981, Velando et al. 2006; Vale and Little 2012; Brannelly et al. 2016). While more study is needed to conclusively establish a positive effect of *Saprolegnia* on *E. baikalensis* reproduction, the *E. baikalensis-Saprolegnia* interaction may represent an interesting system for the study of host-parasite evolution.

### Modeling results and comparison to long-term field data

Our population model reproduced some aspects of field observations on densities of *E. baikalensis* and *Saprolegnia* infection prevalence, providing partial support for our hypothesis that warmer temperatures will have negative population-level impacts on *E. baikalensis* (H3, H4). There were, however, also important differences between the predictions of our model and field data that contrast with our hypotheses H3 and H4. While all formulations of our model predict lower densities of pelagic *E. baikalensis* populations under warmer conditions, the long-term data show no significant differences in the density of adult *E. baikalensis* between years of different temperatures and a possible positive effect of temperature on juvenile densities (Figs. 6, 7). This mismatch may be due to the way we modeled DVM behaviour of *E. baikalensis*. In our model, all *E. baikalensis* performed DVM, spending half of the day in the hypolimnion and half of the day in the epilimnion, regardless of epilimnetic temperatures. In the model, DVM was shown to provide a “thermal refuge” for *E. baikalensis*, allowing it to escape some of the negative effects of warming and persist even under warm conditions (supporting hypothesis H5). Detailed studies of DVM in Lake Baikal (Afanasyeva 1977) and other systems (e.g., Haney 1988; Hays et al. 2001) show that DVM behaviour of copepods is much more nuanced than simulated in our model, with copepod DVM responding to various cues, including food, light, temperature, and predators and also differing for different life stages. In Lake Baikal, Afanasyeva (1977) showed that *E. baikalensis* tended to avoid migrating into the upper part of the epilimnion (0–5 m) when its water temperature exceeded 11–14°C.

The hypothesis that zooplankton modify their DVM behaviour to maximize fitness while balancing different environmental pressures (including predation, food availability, damage from UV radiation and thermal stress) has received wide support (Haney 1988; Loose and Dawidowicz 1994; Williamson et al. 2011). It appears that *E. baikalensis* in Lake Baikal may also adjust their DVM behaviour in response to temperature. The positive relationship between *E. baikalensis* juvenile densities and surface temperature may be a function of such fine-tuning of DVM behaviour, where in warm years *E. baikalensis* have a wider range of thermal regimes in which to maximize feeding, growth, gonad maturation, reproduction and longevity. It is also possible that other factors, for which we did not account in our model, are responsible for a lack of a negative effect of warm temperatures on *E. baikalensis*. For example, primary production rates may be higher during warm years providing more food to *E. baikalensis* and compensating for the negative effects of temperature; Yoshida et al. (2003) found higher primary production rates in the surface water of Lake Baikal in warmer periods of the year and Izmest’yeva et al. (2011) showed a positive correlation between surface temperature and chlorophyll concentrations using long term monitoring data (1979-2002). Reduced cropping of *E. baikalensis* by cold-adapted invertebrate (the pelagic amphipod *Macrohectopus branickii* Dyb.) and vertebrate (various species of endemic fish, mainly *Comephorus baicalensis* Pallas, *C. dybowskii* Kototneff, *Cottocomephorus grewihgki* Dybowski, *C. inermis* Jakowlew) predators of Lake Baikal represents another possible explanation for the mismatch between model predictions and observations. Nonetheless, both our model and field observations indicate that the ability to perform (and possibly fine tune) DVM behaviour provides an important thermal refuge, potentially enabling *E. baikalensis* to persist in the pelagic region of the lake in the face of moderate warming.

In addition to providing a refuge from the direct negative effects of warm temperatures, DVM may also provide *E. baikalensis* with a partial refuge from impacts of *Saprolegnia* infection. Regulation of body temperature by poikilothermic hosts to slow the growth of parasites has been demonstrated in insects (Müller and Schmid-Hempel 1993; Inglis et al. 1996) and modeling suggests that it may be important in our study system. *Saprolegnia* growth and lethality increase with temperature, and the ability to spend a portion of the day in the cold hypolimnion may slow the progress of disease in infected individuals. This finding contrasts with some studies on parasites of *Daphnia* in shallow lakes, where DVM may actually increase infection prevalence by increasing the spatial overlap between hosts and parasites spores, which are concentrated in the hypolimnion (Decaestecker et al. 2002). Thus, the role of DVM as a refuge from warm-loving parasites likely depends on both the individual thermal tolerances of the host and parasite as well as the route of parasite transmission in the system, possibly being most important in deep lakes or the ocean.

### Population-level significance of *Saprolegnia* infection

Reports from the Russian-language literature (Kozhov 1963, Afanasyeva 1977) and long-term monitoring data suggest that, at least in some years, *Saprolegnia* can infect and kill a large fraction of the pelagic *E. baikalensis* population. Despite the potential ecological importance of *Saprolegnia* in the pelagic ecosystem of Lake Baikal, many questions about the impacts of *Saprolegnia* on *E. baikalensis* are unanswered. For example, it is unclear how frequently outbreaks occur. As mentioned earlier, we are unsure whether *Saprolegnia* presence was recorded systematically in the long-term monitoring data or only during some outbreak years. It is also unclear what triggers *Saprolegnia* outbreaks. While some authors hypothesized a link to warm temperatures (Kozhov 1963; Afanasyeva 1977), the years in which *Saprolegnia* presence was recorded in the long-term data were not unusually warm. The spatial extent and dynamics of *Saprolegnia* epizootics are also entirely unexplored and are impossible to evaluate with the long-term data, since these data come from only one sampling station.

Another important knowledge gap concerns the transmission of *Saprolegnia* among *E. baikalensis*. Oomycete infections are transmitted by motile zoospores with limited lifespans (Walker and Van West 2007). In infected *E. baikalensis*, zoospores are not released until days after the death of infected individuals, by which point many of the infected individuals sink out of the upper water layers where most susceptible hosts are concentrated (Fig. 5). This suggests that epizootics may be likelier to start during turbulent conditions when dead individuals remain in the upper layers longer or when zoospores are mixed into the upper water layers (storms or deep convective mixing periods), or in shallow areas of the lake where contact between infectious dead and susceptible individuals is maximized. Transport of spores of the parasitic yeast *Metschnikowia* from nearshore areas by storms and density currents have been hypothesized to play a role in initiating epizootics in *Daphnia* (Cáceres et al. 2006; Hall et al. 2010), and such mechanisms may be also important for *Saprolegnia* epizootics in Lake Baikal.

Among the most challenging aspects of modeling epizootics is estimation of the transmission rate parameter (β), which includes the movement of parasites among hosts as well as the probability of infection once a host is encountered. Estimating β is a non-trivial task even in well-studied host-parasite systems (Kirkeby et al. 2017). We had very little information to guide parameterization of β or assess whether β is temperature-dependent. Given this uncertainty, we sought to explore the consequences of varying β and its temperature dependence for *E. baikalensis* for its heuristic, rather than strictly predictive value.

Research on host-parasite interactions in aquatic systems often predicts higher incidence of infection with warmer temperatures (Marcogliese 2008; Karvonen et al. 2010). However, this is not always the case; for example, Hall et al. (2006), showed that interactions between the thermal optima of parasites and their hosts (as well as of host’s predators) can result in diverse temperature-infection prevalence relationships. We found that the shape of the β - temperature relationship had large and opposite effects on predicted infection prevalence under different temperature scenarios (Fig. 7). In a temperature-independent β scenario, the highest rates of infection and greatest negative impacts of *Saprolegnia* were predicted under the coldest temperature scenarios. This result is a function of the higher densities of susceptible individuals under colder scenarios and density-dependent transmission of *Saprolegnia* in the model. Interestingly, the highest incidence of infection was still predicted for the warmer periods of the year, likely because of increased temperature-driven mortality of infected individuals. Allowing β to increase with temperature resulted in a different pattern of modeled infection prevalence relative to the “constant-β” scenario. Both the “mortality rate”-and “growth on agar”-based temperature-β relationships predicted higher incidence of infection for the warmer scenarios; the steeper slope of the temperature-β relationship in the latter scenario resulted in higher incidence of infection across all temperature scenarios relative to the “mortality-rate”-based temperature-β relationship.

Deciding which of our temperature-β scenarios is more realistic, and hence better simulates the infection processes that occur in Lake Baikal, is difficult. We have shown that the time between exposure and death (and hence ability to release infective zoospores) may decrease with increasing temperature. On the other hand, the sinking of dead individuals prior to zoospore release means that spore maturation and host-seeking may occur in the deep and cold hypolimnion, and hence β would be independent of surface temperature. Given the large differences in prediction of different temperature-β scenarios, determining which scenario is more realistic will enable better predictions of how *Saprolegnia* may affect the Lake Baikal ecosystem in the future.

Some of the temperature-dependent β scenarios predicted very high infection prevalence by *Saprolegnia* in the *E. baikalensis* population. Interestingly, infection-prevalence did not have a linear relationship with the negative effect of the infection. This result is likely because of the non-linear relationships between temperature, infection rates and the mortality rates of infected and uninfected adults and juveniles. These findings suggest an interesting possibility: since infection can only be diagnosed after death, it is possible for a large proportion of the *E. baikalensis* population to be infected with *Saprolegnia* without showing strong negative effects under normal temperature conditions, and for the negative impacts to become rapidly manifested during temperature increases. However, we believe this scenario is unlikely to be the norm in Lake Baikal. In dozens of experiments that we conducted with *E. baikalensis* in 2012 and 2013, the development of *Saprolegnia* hyphae after death was observed in only a small number of individuals that were not intentionally exposed to *Saprolegnia* zoospores. Thus, *Saprolegnia* epizootics are probably an episodic occurrence in the lake rather than a regular feature of the pelagic ecosystem. Additional study is needed to determine the frequency, causes and consequences of *Saprolegnia* epizootics in the simple pelagic food web of Lake Baikal.

### Implications in the context of environmental change

Lake Baikal is frequently regarded as a unique ecosystem due to its old age, high volume, and notable biodiversity and endemism, yet most of the threats it now faces are not unique (Hampton et al. 2018). In the vast majority of the world’s lakes for which data are available, significant warming has occurred (O’Reilly et al. 2015). This warming has been suggested to negatively affect the endemic taxa that have evolved in both the warmest and the coldest of these ecosystems (Hampton et al. 2018). In the relatively hot African Rift Lakes that harbor extraordinary endemic biodiversity, fish and other endemic organisms may already be living near the limits of their thermal tolerances (O’Reilly et al. 2015) and it is unclear how long the endemic stenotherms of cold systems can be sustained in thermal refugia (Moore et al. 2009).

Surface temperatures of Lake Baikal have been rising for decades (Shimaraev and Domysheva 2013). Piccolroaz and Toffolon (2018) predict a mean and maximum increase in summer surface temperatures of 1.9 and 4°C for Lake Baikal by the middle of the 21^st^ century. Our model and long-term monitoring data suggests that the negative effects of this temperature increase may be moderated by the ability of *E. baikalensis* to fine-tune their DVM behaviour. However, the direct negative effects of temperature on *E. baikalensis* will likely interact with other aspects of their environment. Besides potential for increased parasitism by *Saprolegnia*, interactions with cosmopolitan species of warm-loving competitors *(Daphnia)* and predators (the cyclopoid copepod *Cyclops kolensis)* – the densities of which have been increasing in recent decades (Hampton et al. 2008; Izmest’eva et al. 2016; Silow et al. 2016) – may have negative consequences for *E. baikalensis*.

In summary, our experimental, field and modelling results show a large thermal mismatch between cold-adapted freshwater copepods and their oomycete parasites in Lake Baikal. The negative effects of parasites are shown to vary seasonally and potentially between cold and warm years as the importance of the hot-parasite thermal mismatch varies. Modelling suggests that DVM by *E. baikalensis* may provide not only more optimal temperature for survival and reproduction but also a potential refuge from the negative effects of parasites. This hypolimnetic refuge may allow populations of *E. baikalensis* and other cold-loving copepods in lakes and oceans to persist even while surface temperatures rise above the optimum range for these organisms. However, temperature-mediated changes in behavior as well as the presence of asymmetric temperature responses for closely interacting species, such as those considered here, are very likely to alter interaction strengths in food webs with unknown impacts on aquatic resources.

## Acknowledgements

This work was supported by National Science Foundation Dimensions of Biodiversity awards DEB-1136637, DEB-1136667, and DEB-1136657. The long-term Lake Baikal data are part of a data sets No. 2005620028 and No. 2014621482 registered with the government of the Russian Federation. E.V. Pislegina, E.A. Silow and M.A. Timofeyev were supported by a Ministry of Higher Education and Research of Russian Federation award #6.1387.2017 and by funding from the Lake Baikal Foundation (https://baikalfoundation.ru/project/tochka-1/). We are grateful to Steve Katz, Ed Theriot, Elena Litchman, Lev Yampolsky, Stephanie Labou and Meghan Duffy for useful advice that shaped the research.

