## Supplemental text, table and figures for "Hot and sick: impacts of warming and oomycete parasite infection on endemic dominant zooplankter of Lake Baikal"

**Supplementary information:**

Phylogenetic identification of *Saprolegnia*

Phylogenetic identification of *Saprolegnia* was accomplished using analysis of COI and ITS genes. DNA was isolated from “mycelia” grown on agar by bead-beating with glass beads in a solution containing lysis buffer and RNAse. The Plant DNAEasy (Qiagen) protocol was used to purify DNA from this lysate. The mitochondrial cytochrome oxidase I (COI) gene and internal transcribed spacer (ITS) of the nuclear ribosomal operon were PCR-amplified using primers from Robideau et al. (2011). For COI, we used the OomCoxI-Levup (5’-TCAWCWMGATGGCTTTTTTCAAC-3’) and Fm85mod (5’-RRHWACKTGACTDATRATACCAAA-3’) primer pair, both for PCR amplification and Sanger sequencing. For ITS, we first amplified a region using the primer pair: UN-up18S42 (5’-CGTAACAAGGTTTCCGTAGGTGAAC-3’) and UN-lo28S1220 (5’-GTTGTTACACACTCCTTAGCGGAT-3’). This stretch encompassed the ITS region and a part of the downstream 28S gene. To sequence only the target ITS fragment, we substituted UN-lo28S1220 with UN-lo28S22 (5’-GTTTCTTTTCCTCCGCTTATTGATATG-3’) in the sequencing reaction (Robideau et al. 2011). The resulting sequences were edited to remove poor quality base calls and then submitted to Genbank (accession numbers SAP1, COI: MN058729; SAP1, ITS:  MN046970).

For phylogenetic analysis, our newly generated sequences were combined with previously published data available in Genbank. For COI, we downloaded a total of 43 *Saprolegnia* sequences and aligned them by hand, guided by their conceptual amino acid translations. For ITS, a bulk download from Genbank resulted in several hundred *Saprolegnia* sequences, many of which were taxonomically redundant (i.e., same species) or non-redundant but identical (i.e., different species but same sequence). This set was thinned by finding the ITS data for *Saprolegnia* strains with available COI sequence arriving at a total of 42 Genbank sequences. We aligned the ITS data using MAFFT v7.273 (Katoh and Standley, 2013) with the option ‘--auto’ which lets the program choose the most suitable alignment method. The resulting alignments of COI (44 sequences, 680 base pairs, bp) and ITS (43 sequences, 718 bp) were then analyzed independently and jointly in a concatenated COI+ITS alignment (44 sequences, 1398 bp).

Phylogenetic trees were inferred using IQtree v1.6.3 (Nguyen et al. 2015). We analyzed the single-gene alignments unpartitioned, but allowed 4 partitions for the longer concatenated alignment: 3 for each codon position of the COI gene and 1 for the entire ITS marker. For each alignment, we first ran a model-selection routine to identify the best model of nucleotide substitution and rate variation across alignment columns. For the concatenated alignment, we also ran a partition merging procedure that identified whether any pair of partitions could be merged without a significant loss in model fit. After identifying the best models and partitioning scheme, we ran five independent maximum likelihood (ML) optimizations for the single-gene alignments and 25 for the concatenated alignment. We ran IQtree with default settings and assessed node support using the Shimodaira-Hasegawa-like approximate Likelihood Ratio Test (SH-aLRT; Guindon et al. 2010) and the UltraFast Bootstrap approximation (UFBoot; Minh et al. 2013) with 1000 pseudoreplicates for each. In addition to the IQtree searches, we used RAxML v8.1.7 (Stamatakis, 2014) to repeat the analysis of the concatenated COI+ITS alignment. We ran 25 RAxML searches with a GTR+Г4 model of nucleotide substitution for each partition. We assessed node support with RAxML’s Rapid Bootstrap method (Stamatakis et al. 2008) and 1000 pseudoreplicates. In all cases, we treated the tree with highest log-likelihood as the ‘best tree’, and we rooted the trees by setting *Saprolegnia asterophora* as sister to all other taxa.

In the COI phylogeny, the *Saprolegnia* strain isolated from *E. baikalensis* was reconstructed as a sister lineage to a large clade of *Saprolegnia* species including *S. diclina, S. parasitica, S. ferax, S. unispora* and others (SH-aLRT/UFBoot = 93.4/90). In the ITS tree, the *Saprolegnia* strain isolated from *Epischura* was a strongly supported sister lineage to a clade of two isolates of *S. diclina* (SH-aLRT/UFBoot = 96.4/99). This result also held for the analysis of the combined COI+ITS data (Fig. S3), where both IQtree and RaxML found strong support for a sister relationship between the *E. baikalensis*-infesting *Saprolegnia* and *S. diclina* (IQtree: SH-aLRT/UFBoot = 93.5/99; RAxML Rapid bootstrap=98%). Although it might appear that the *E. baikalensis*-infesting *Saprolegnia* is most closely related to *S. diclina*, strains of several *Saprolegnia* species, including *S. diclina,* were scattered across the tree (Fig. S3), suggesting a mismatch between phylogenetic relationships and species boundaries (i.e., strains of the same species did not form a monophyletic group). Moreover, the lineage to which the *E. baikalensis* -infesting *Saprolegnia* belongs has many short branches implying relatively recent divergences. It is therefore difficult to determine the exact taxonomic identity of our *E. baikalensis*-infesting *Saprolegnia*, but the analysis confirms that it is related to other taxa implicated as parasites of aquatic organisms.

Model:

To examine the population-level effects of seasonal water temperature and *Saprolegnia* infection on *E. baikalensis*, we constructed a population model for *E. baikalensis* using the STELLA modeling environment (Fig. S2). The model used the fourth order Runge-Kutta method to approximate differential equations with a time step of 0.125 days. The model included healthy (uninfected) adults (A_H_), infected adults (A_I_), healthy juveniles (J_H_) and infected juveniles (J_I_). Densities were modeled as:

(1) A_H(t)_ = gJ_H(t-1)_ - m_AH_A_H(t-1)_ - β(A_H(t-1)_ + J_H(t-1)_)(m_AI_A_I(t-1)_ + m_JI_J_I(t-1)_)

(2) A_I(t)_ = β(A_H(t-1)_ + J_H(t-1)_)(m_AI_A_I(t-1)_ + m_JI_J_I(t-1)_) + gJ_I(t-1)_ - m_AI_A_I(t-1)_

(3) J_H(t)_ = r_H_A_H(t-1)_ + r_I_A_I_ - gJ_H(t-1)_ - m_JS_J_H(t-1)_ - β(A_H(t-1)_ + J_H(t-1)_)(A_I(t-1)_ + J_I(t-1)_)

(4) J_I(t)_ = β(A_H(t-1)_ + J_H(t-1)_)(A_I(t-1)_ + J_I(t-1)_) - gJ_I(t-1)_ - m_JI_J_I(t-1)_

The model was parameterized using a combination of experimentally-determined and literature-estimated values for life stage- and infection status-specific: mortality rate (m), reproductive rate (r), maturation rate (g), and infection transmission rates (β) (Table 1). The temperature dependences of various rates were determined for 5, 10, 15 and 20°C and linear interpolation was used to estimate rates between these temperatures; 20°C rates were used for all temperatures above 20°C. Individuals in the model had an experimentally-derived, temperature-, life stage- and infection status-specific mortality rates (m). Healthy individuals could become infected through “mass-action” transmission (McCallum et al. 2001), which depends on the density of healthy adults and juveniles and dead infected individuals (since zoospores are released only after death) that determine the encounter rate at each time step. The transmission rate (β) includes both the encounter rate plus the probability of infection after encounter. Infection rates for adults and juveniles are assumed to be the same. Adults have experimentally-determined, temperature and infection status-specific reproductive rates (r), with both susceptible and infected adults producing only healthy juveniles (*Saprolegnia* is transmitted horizontally; Kar 2016). Juveniles have temperature-dependent, literature-estimated growth (maturation) rates (g).

We used long-term temperature data (1948-2002) from 0–25 m depths at a pelagic station in Lake Baikal (recorded approximately every 2 weeks) to construct seasonal temperature scenarios representative of ‘cold’, ‘cool’, ‘average’, ‘warm’ and ‘hot’ years in the pelagic zone (same definitions of temperature categories as in the long-term data analysis). We fit generalized additive models (GAMs, R package ‘mgcv’) to long-term temperature data within each temperature category to estimate daily-step temperatures through the stratified period of the year (period when surface temperatures are above 4°C); daily time-step temperatures were extracted from the GAM models using the *predict()* function in R. We also used seasonal temperature data from a shallow embayment of the lake (Proval Bay; Kozhov 1963) to simulate conditions in a warm shallow bay. We parameterized a temperature scenario for a large bay (e.g., Barguzin, Chivyrkuy or warm parts of Maloe More) by averaging the temperature curves for Proval Bay and the “average” pelagic scenario. To examine the role of DVM on *E. baikalensis* populations under different temperature and infection scenarios, we created models where *E. baikalensis* spent half of the day in the hypolimnion (by modeling daily temperature as an average of hypolimnetic and epilimnetic temperatures) and models where *E. baikalensis* were restricted to the epilimnion (as might be the case in a shallow region of the lake).

In the model, initial densities of healthy adult and juvenile *E. baikalensis* (at DOY 1; Jan. 1) were set equal to average densities of adults and juveniles at DOY 355-10 (Dec. 21–Jan. 10) across all years in the long-term dataset. Initial density of infected individuals was set as ~1% of the total population based on average densities of infected (dead) individuals for December and January in the long-term data for years for which *Saprolegnia* was recorded.

Maturation rates, estimated from Afanasyeva (1977), were 0.0056 for 5°C and 0.011 for 10°C and higher. The model used experimentally determined reproductive rates for the different temperatures. Although differences in nauplii production rates between egg sacks of *Saprolegnia*-exposed and control individuals occurred (see results), we averaged reproductive rates across all individuals in the model. The main reason for this averaging was that we are not certain of the mechanism that could underlie the observed increase in nauplii production in *Saprolegnia*-exposed individuals (see results section) or how to properly model it (i.e., whether egg sacks need to be modeled as a separate compartment that can become infected or whether the reproductive “advantage” of *Saprolegnia* infection is conferred from the parent becoming infected). While this is a possible shortcoming of our model, we believe that the model still provides a useful tool for exploration of the dynamics of *E. baikalensis* given the many unknowns of this host-parasite system.

Mortality rates were determined from a combination of experimental results and trial modeling. Recognizing that our lab-determined mortality rates are likely not representative of rates in the lake, we created a simplified version of our population model with a constant temperature of 5°C and ran it for 175 days, corresponding to the time between the start of the calendar year (DOY 1, Jan. 1) and the typical date when surface temperatures exceed 4°C in Lake Baikal (DOY 175, Jun. 24). We experimented with different values of juvenile and adult mortality rates until we found rates that resulted in population dynamics that corresponded to the gradual increase in densities observed in the long term data between DOY 0 and day 175 (see results section). We used these rates as the ≤5°C “baseline” rates and established the mortality rates for 10, 15, and 20°C based on the relative difference in mortality rates between these temperatures in the lab experiments.

The infection rate parameter (β) is notoriously difficult to determine, even for well-studied systems, and the choice of β can have large effects on model outcomes (Kirkeby et al. 2017). Given that the *E. baikalensi*s *– Saprolegnia* host-parasite relationship has received almost no study, the choice of the value for β used in the model was difficult, as was determining whether to model β as a temperature-dependent or temperature-independent parameter (Lively et al. 2014). We therefore compared the output of model scenarios parameterized with three different β-temperature functional relationships. First, we determined a “baseline” β by using different values of β in the constant 5°C, 175-day model used to determine baseline mortality rates. Our goal was to find a value for β that produced a similar daily number of deaths of infected individuals to that observed in the long-term data for years where *Saprolegnia* was observed in the plankton. Actually, since the number of infected individuals observed in the long-term plankton data on each particular sampling occasion likely represents approximately 5 days-worth of mortality (since it takes ca. 1 day for *Saprolegnia* hyphae to become visible after death and assuming an ca. 5 day persistence of dead individuals in the upper 500 m of the water column; Kirillin et al. 2012), we sought a β value that produced daily mortality rates of infected individuals corresponding to 1/5 of the average number of recorded infected individuals in the long-term data at the end of the spring mixing season across all years. This baseline value was estimated as 0.0008. We then parameterized 3 scenarios modeling different temperature–β relationships. The first scenario was the temperature invariant scenario, where β was 0.0008 individuals/m^2^/day throughout the year. The second scenario, called the “mortality” scenario, used β of 0.0008 for temperatures ≤5°C and the experimentally-determined difference in mortality rates of infected and uninfected *E. baikalensis* (averaged for adults and juveniles) at experimental temperatures to estimate the temperature-virulence relationship for *Saprolegnia*. The values of β for 10, 15 and 20°C were then estimated based on the relative differences in *Saprolegnia*-caused mortality at the different temperatures. The third scenario, called the “agar” scenario, also used a β value of 0.0008 for ≤5°C; values for 10, 15 and 20°C were estimated relative to the 5°C baseline from the growth rates of *Saprolegnia* on agar.

Since our primary interest was to explore the effect of temperature variation on *E. baikalensis* populations, we ran our full models only for the duration of the stratified summer season, when surface temperatures were >4°C. This period differed for different temperature scenarios, ranging from DOY 170–330 (Jun. 19–Nov. 27) in the “cold pelagic” scenario to DOY 129-303 (May 9–Nov. 1) in the “Proval Bay” (warm water) scenario.

**Supplementary tables:**

Table S1: *Epischura baikalensis* average times when last seen alive in an experiment (includes individuals that died and survived till end of experiment) and predicted survival times from parametric survival models

| Group | Mean survival time (±SD), days | Predicted survival time, days |
| --- | --- | --- |
| *Epischura* adults |  |  |
| 5°C, control | 22.4±1.35 | 215.00 |
| 5°C, Saprolegnia | 21.94±2.49 | 175.50 |
| 10°C, control | 21.31±4.08 | 127.88 |
| 10°C, Saprolegnia | 18.44±5.75 | 40.23 |
| 15°C, control | 15.16±9.06 | 23.76 |
| 15°C, Saprolegnia | 11.53±9.15 | 15.37 |
| 20°C, control | 3.91±3.58 | 3.91 |
| 20°C, Saprolegnia | 1.5±1.35 | 1.50 |
| *Epischura* nauplii |  |  |
| 5°C, control | 16.8±7.15 | 24.00 |
| 5°C, Saprolegnia | 15.2±6.85 | 20.91 |
| 10°C, control | 14.08±4.21 | 15.57 |
| 10°C, Saprolegnia | 13.56±4.29 | 14.81 |
| 15°C, control | 7.29±3.47 | 7.83 |
| 15°C, Saprolegnia | 5.66±3.55 | 6.50 |
| 20°C, control | 3.15±1.48 | 3.49 |
| 20°C, Saprolegnia | 2.07±1.24 | 2.41 |

Table S2: Summary (average ± SD) of *Epischura baikalensis* reproductive parameters in the control group (Control) and in the *Saprolegnia* exposure group, separated based on whether egg sacks developed visible hyphae. N/A means no egg sack production or hatching.

|  | Control | Exposed, no hyphae | Exposed, with hyphae |
| --- | --- | --- | --- |
| Time to 1^st^ egg sack (days)  5°C  10°C  15°C  20°C | 5.94 ± 2.19  4.87 ± 2.49  3.58 ± 1.40  N/A | 6.02 ± 2.10  5.15 ± 0.98  4.31 ± 2.12  N/A | 5.48 ± 2.89  3.67 ± 1.42  1.82 ± 0.99  N/A |
| Time to hatch (days)  5°C  10°C  15°C  20°C | 12.70 ± 1.33  8.18 ± 0.54  N/A  N/A | 12.56 ± 0.83  8.25 ± 0.45  N/A | 13.15 ± 1.64  8.47 ± 0.50  N/A  N/A |
| Number of nauplii  5°C  10°C  15°C  20°C | 12.18 ± 10.36  7.17 ± 6.06  N/A  N/A | 35.90 ± 10.40  18.23 ± 9.57  N/A  N/A | 24.35 ± 13.69  8.49 ± 7.36  N/A  N/A |

Table S3: Results of GAMM analysis on abundance of *Epischura baikalensis* adults and juveniles (nauplii + copepodites) through a full year and only during the stratified season in years with cold, cool, average, warm and hot summers. Analysis was designed to determine whether there was significant seasonal variation in *Epischura baikalensis* abundance with day of year (DOY) and whether pattern of variation differed between year types (cold, cool, etc.).

| Group | (e)df | F-value | p-value |
| --- | --- | --- | --- |
| Adults, year-round  DOY  Year type | 6.5  4 | 32.2  0.16 | <<0.00001  0.97 |
| Adults, stratified season  DOY  Year type | 1.77  4 | 50.4  0.20 | <<0.00001  0.94 |
| Juveniles, year-round  DOY  Year type | 5.3  4 | 44.9  2.45 | <<0.00001  0.045 |
| Juveniles, stratified season  DOY  Year type | 2.8  4 | 4.25  1.55 | 0.008  0.19 |

**Supplementary figures:**

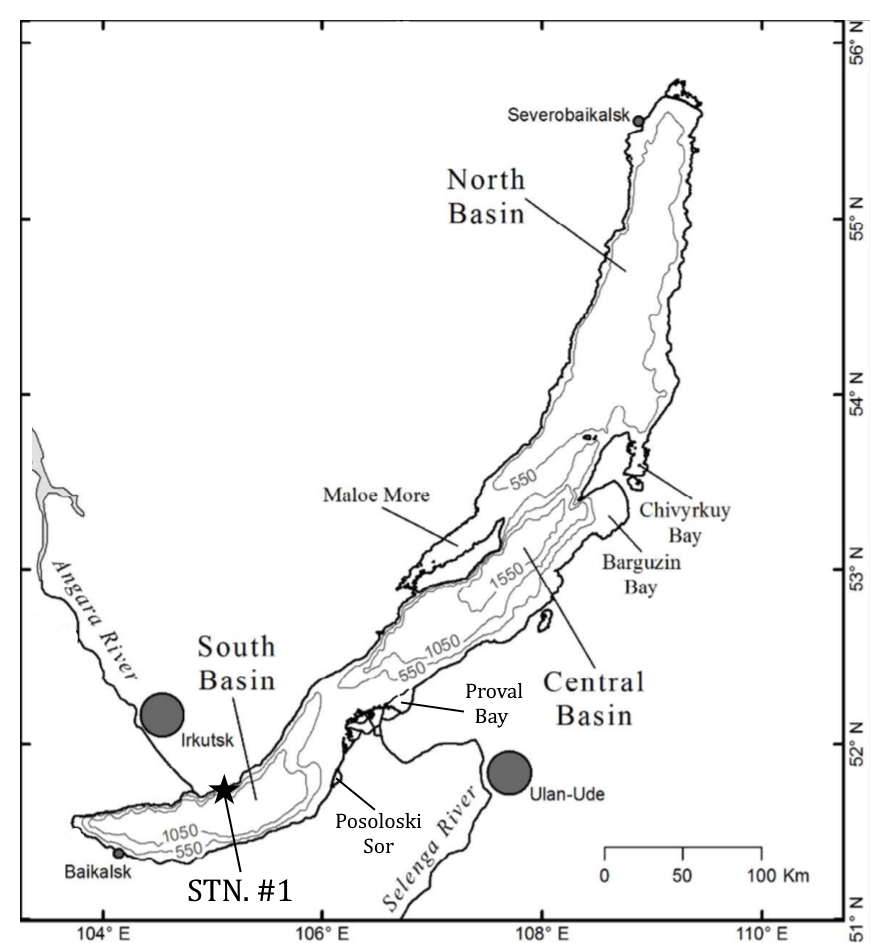

Figure S1: Map of Lake Baikal, showing main human population centers (grey circles), the larger bays and location of Irkutsk State University’s long-term monitoring station (STN. #1). Depth contours are in meters.

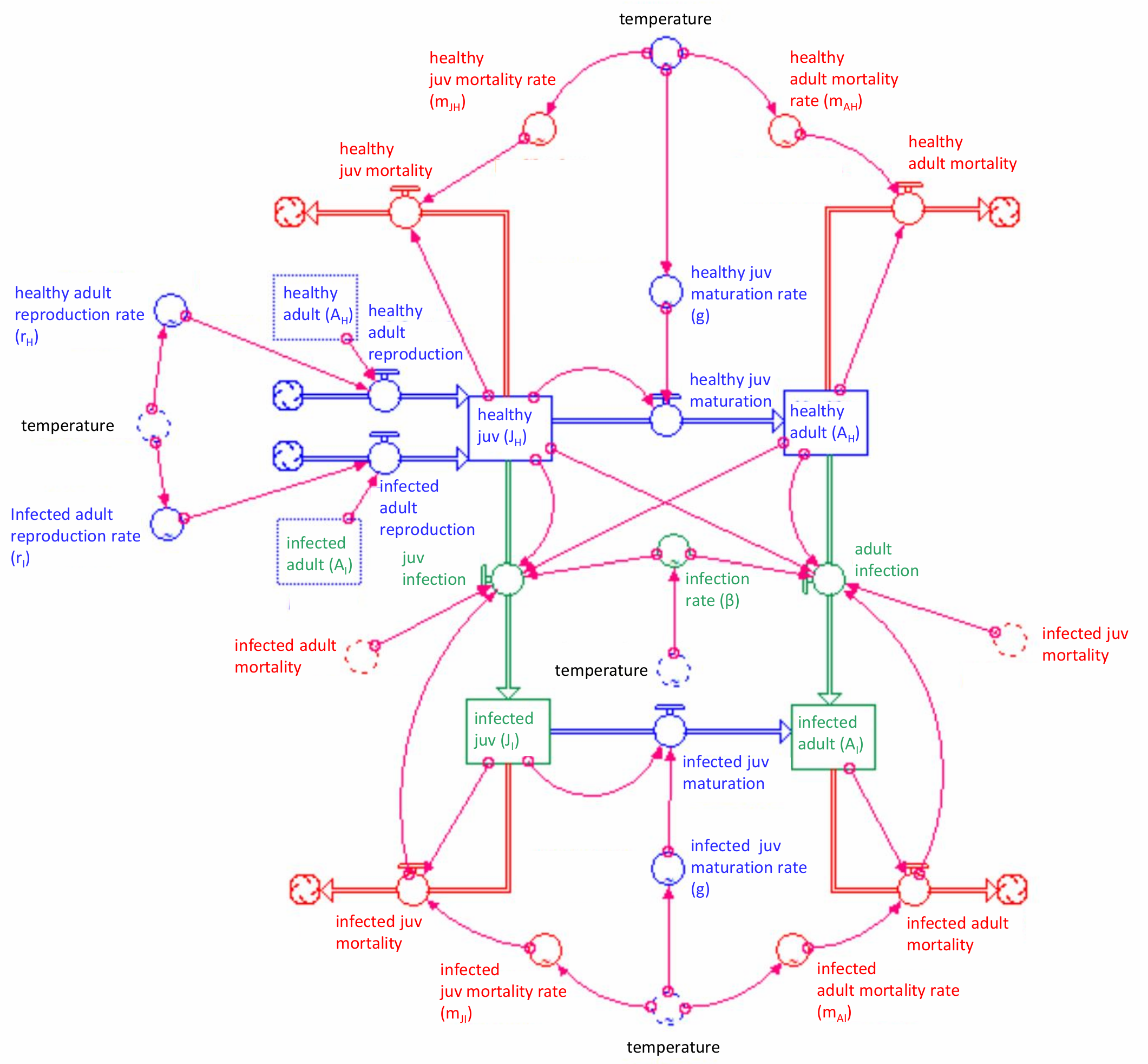

Figure S2: Map of the STELLA model used to model effects of different temperature scenarios and *Saprolegnia* infection on *Epischura baikalensis* populations.

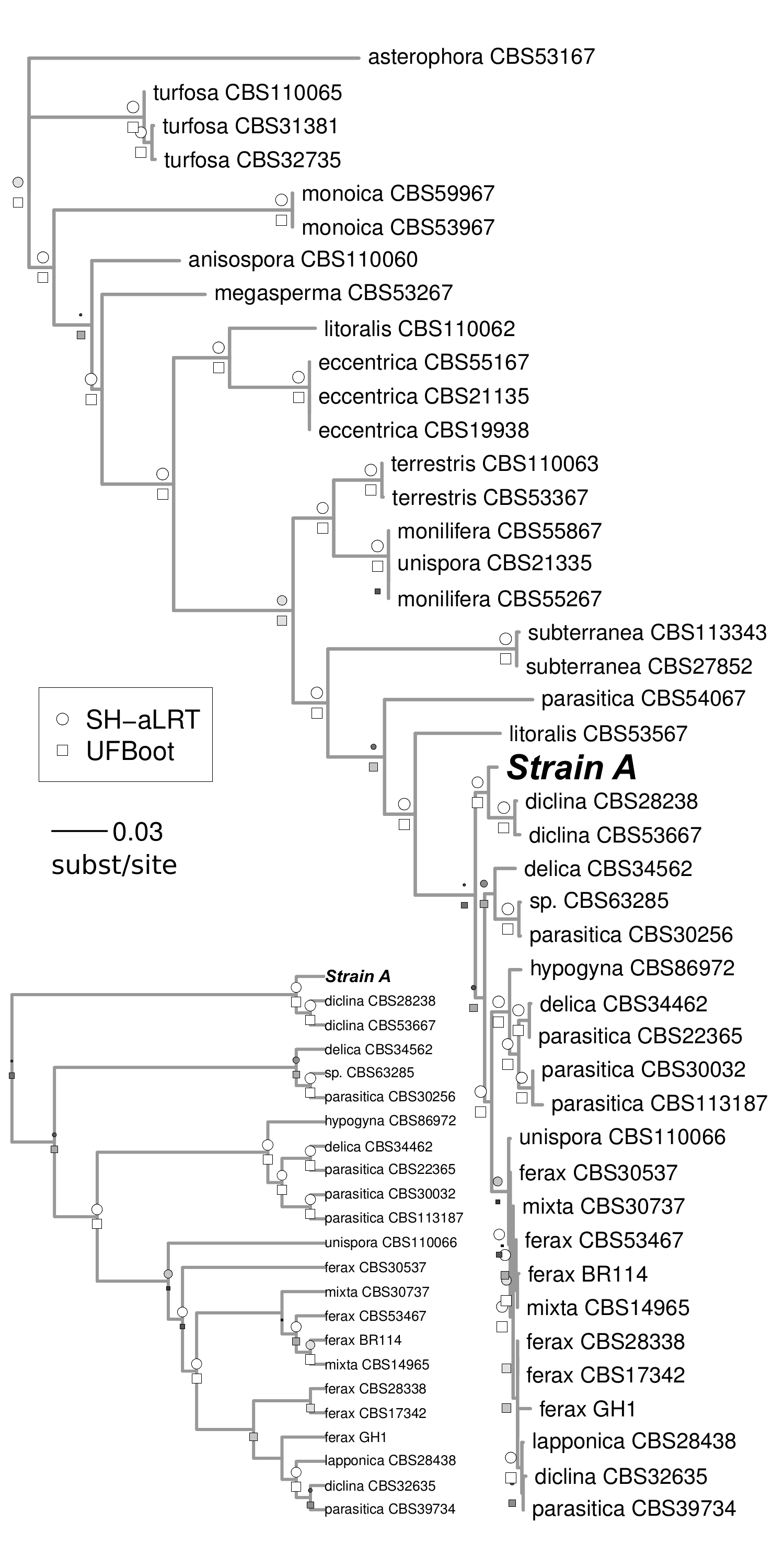

Figure S3: Combined ITS- and COI-based phylogenetic identification of the *Saprolegnia* strain (Strain A) isolated from Lake Baikal *Epischura baikalensis* and used in experiments.

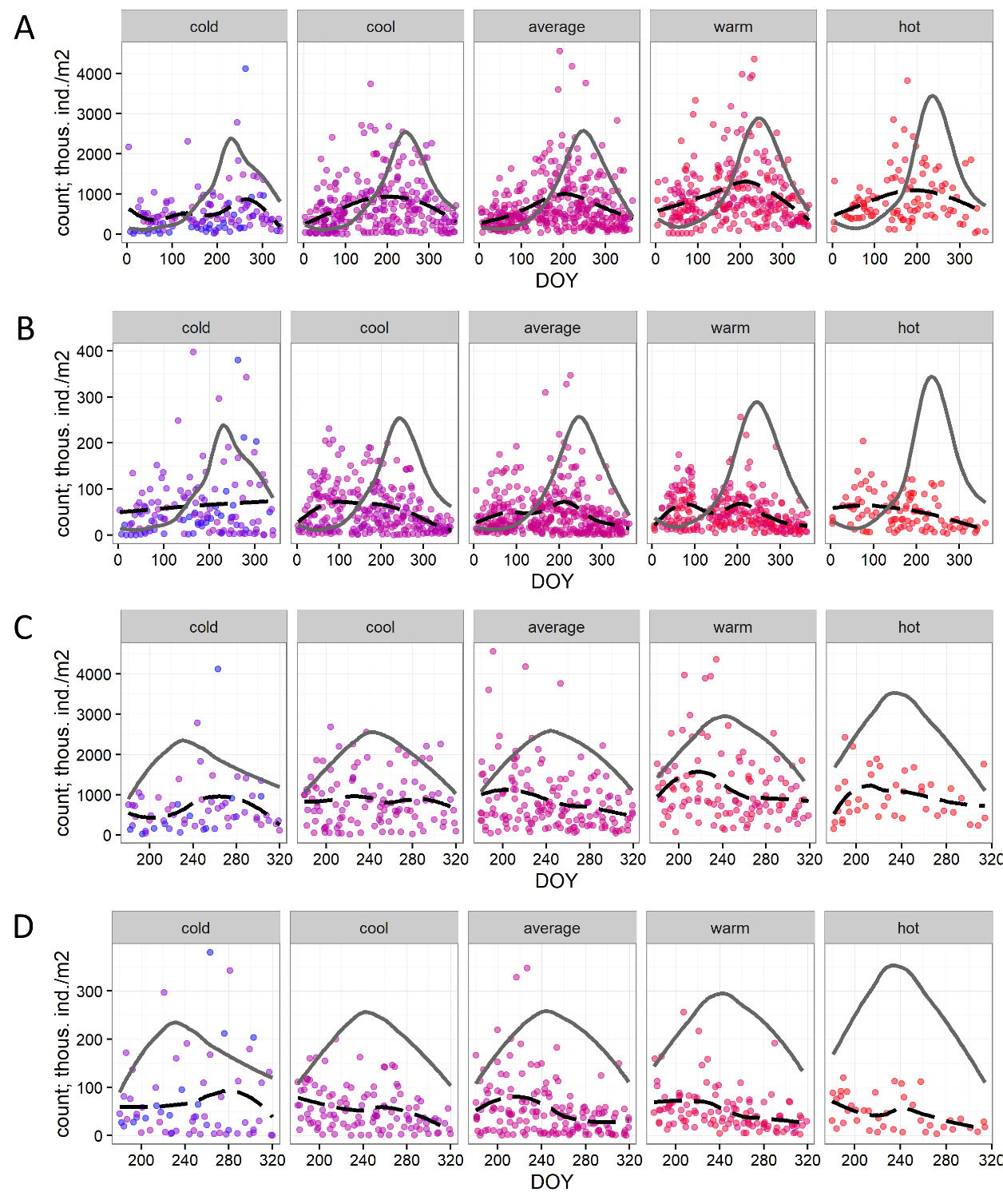

Figure S4: Seasonality of abundance of *Epischura* adults and juveniles (nauplii + copepodites) in the 0-250 m water layer in cold, cool, average, warm and hot years in the pelagic zone of Lake Baikal based on 48 years of data. A) juvenile abundance throughout full year; B) adult abundance throughout full year; C) juvenile abundance throughout stratified period; D) adult abundance throughout stratified period.

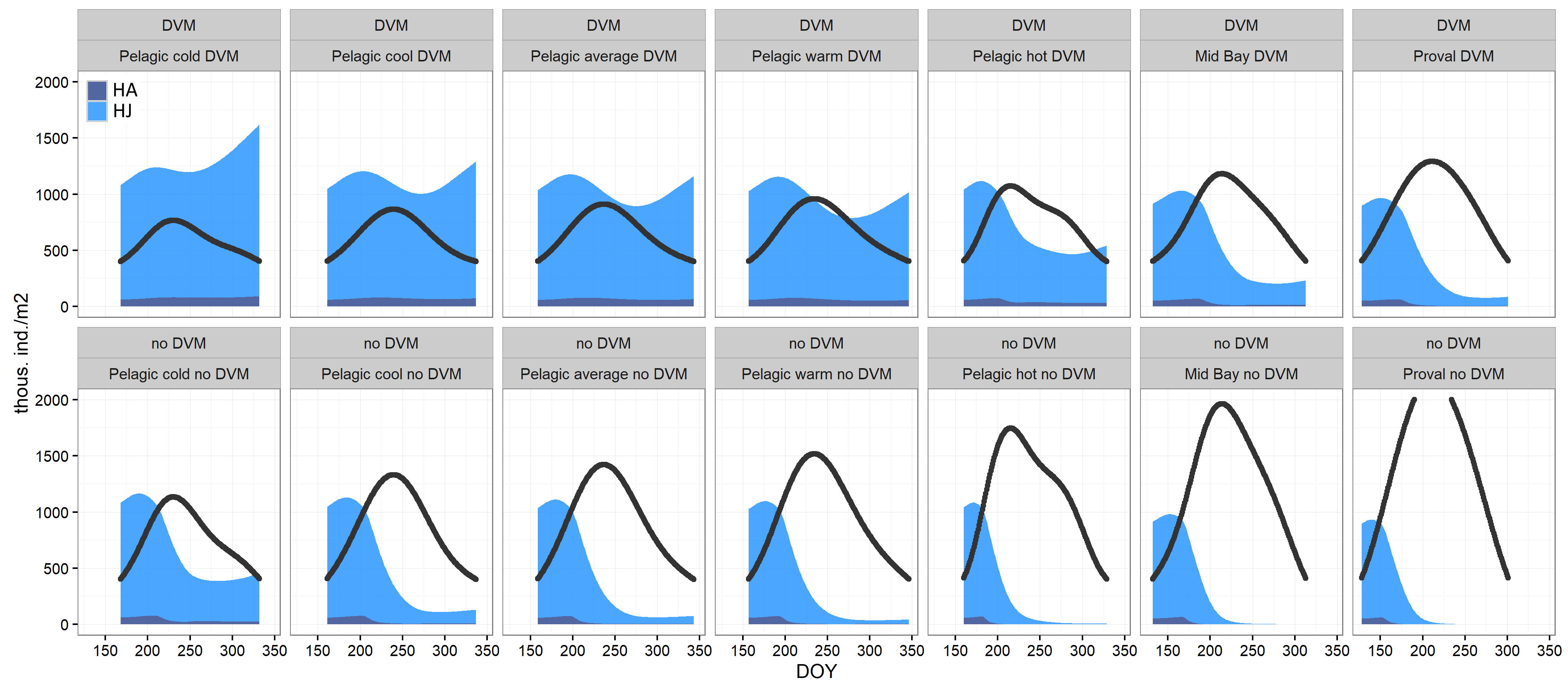

Figure S5: Modeled abundance of *Epischura* during the stratified open water season with no *Saprolegnia* infection. Runs were performed for 7 different temperature scenarios; each temperature scenario was run for theoretical populations performing DVM and theoretical populations not performing DVM. Filled areas correspond to *Epischura* abundance: HA= healthy adults, HJ= healthy juveniles. Black line represents 100*(surface water temperature).

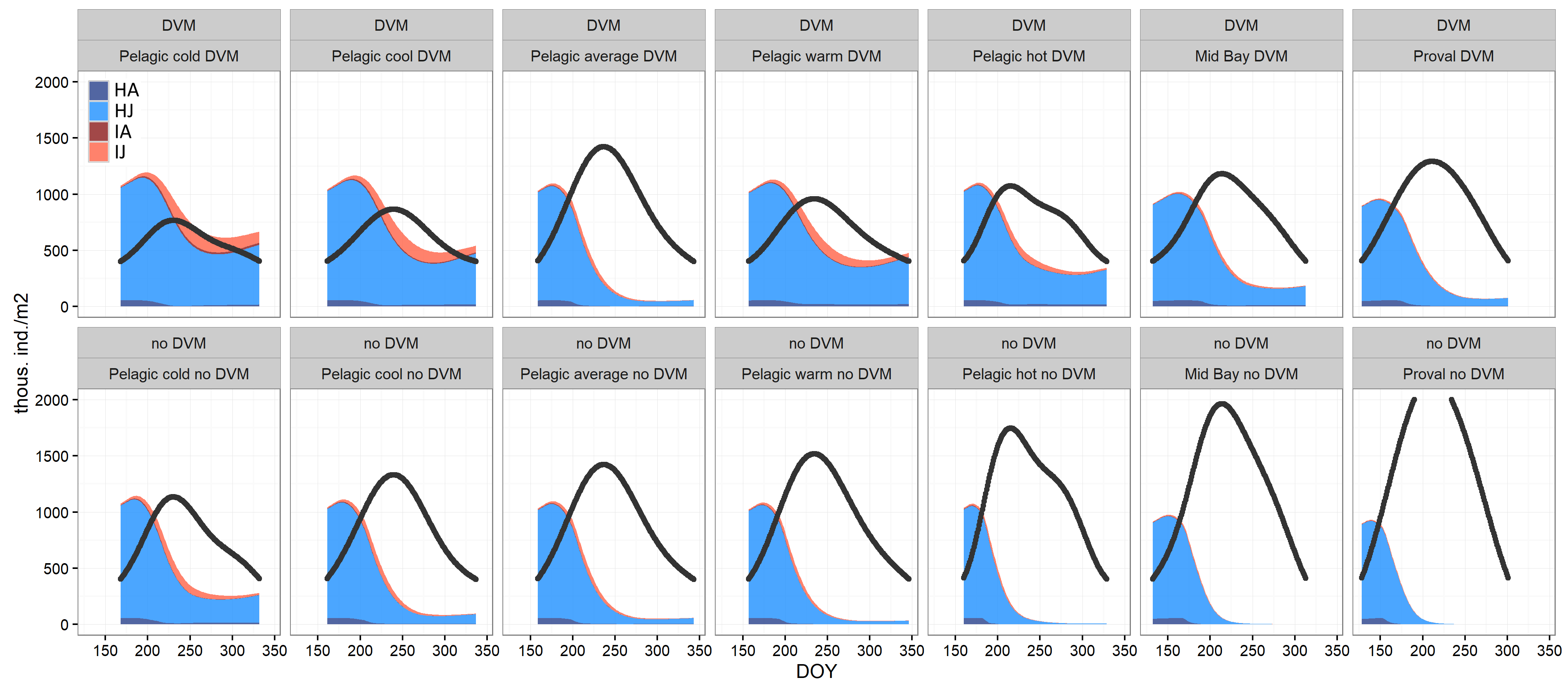

Figure S6: Modeled abundance of *Epischura* during the stratified open water season with constant (temperature independent) *Saprolegnia* transmission rate (β). Runs were performed for 7 different temperature scenarios; each temperature scenario was run for theoretical populations performing DVM and theoretical populations not performing DVM. Filled areas correspond to *Epischura* abundance: HA= healthy adults, HJ= healthy juveniles, IA= infected adults (as thin dark red line below abundance of IJ), IJ= infected juveniles. Black line represents 100*(surface water temperature).

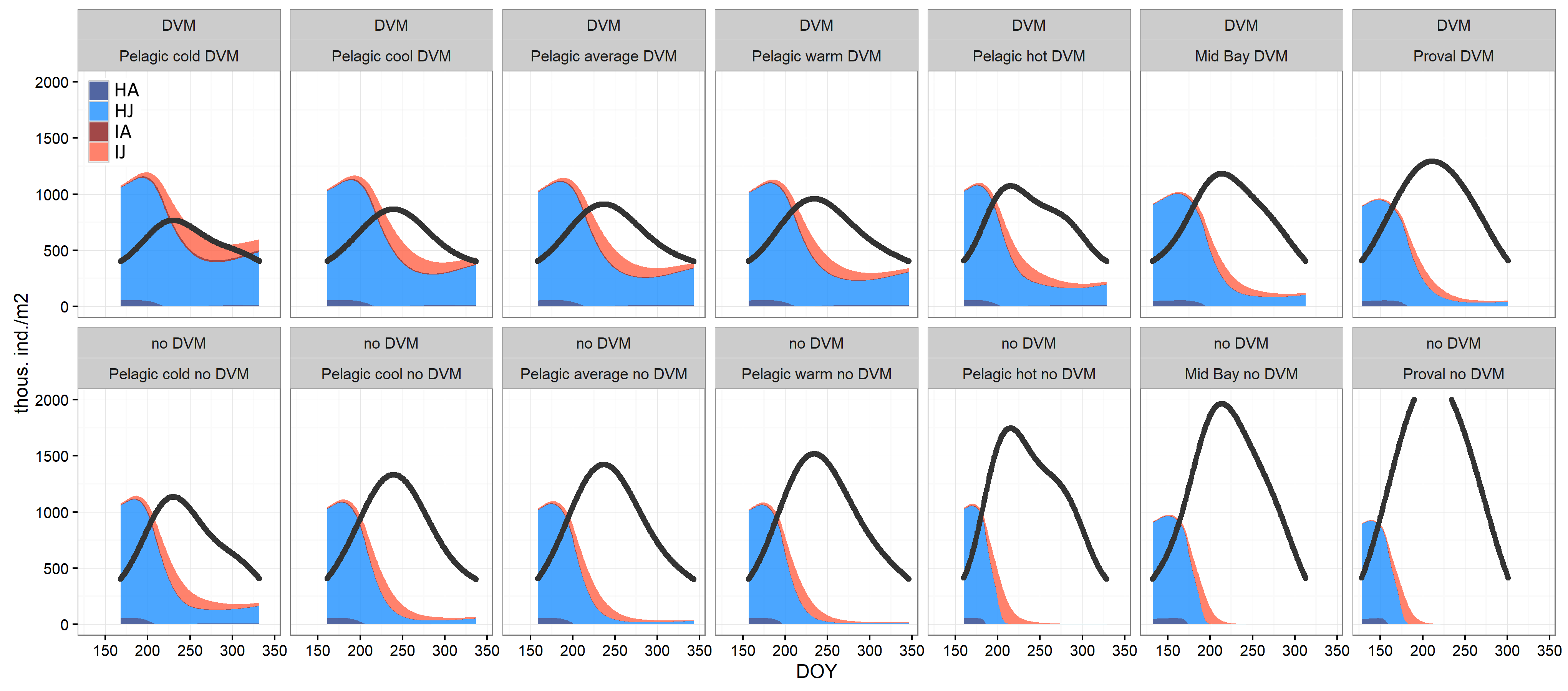

Figure S7: Modeled abundance of *Epischura* during the stratified open water season with temperature-dependent *Saprolegnia* transmission rate (β), modeled based on *Saprolegnia*-infected *Epischura* mortality rates. Runs were performed for 7 different temperature scenarios; each temperature scenario was run for theoretical populations performing DVM and theoretical populations not performing DVM. Filled areas correspond to *Epischura* abundance. HA= healthy adults, HJ= healthy juveniles, IA= infected adults (as thin dark red line below abundance of IJ), IJ= infected juveniles. Black line represents 100*(surface water temperature).

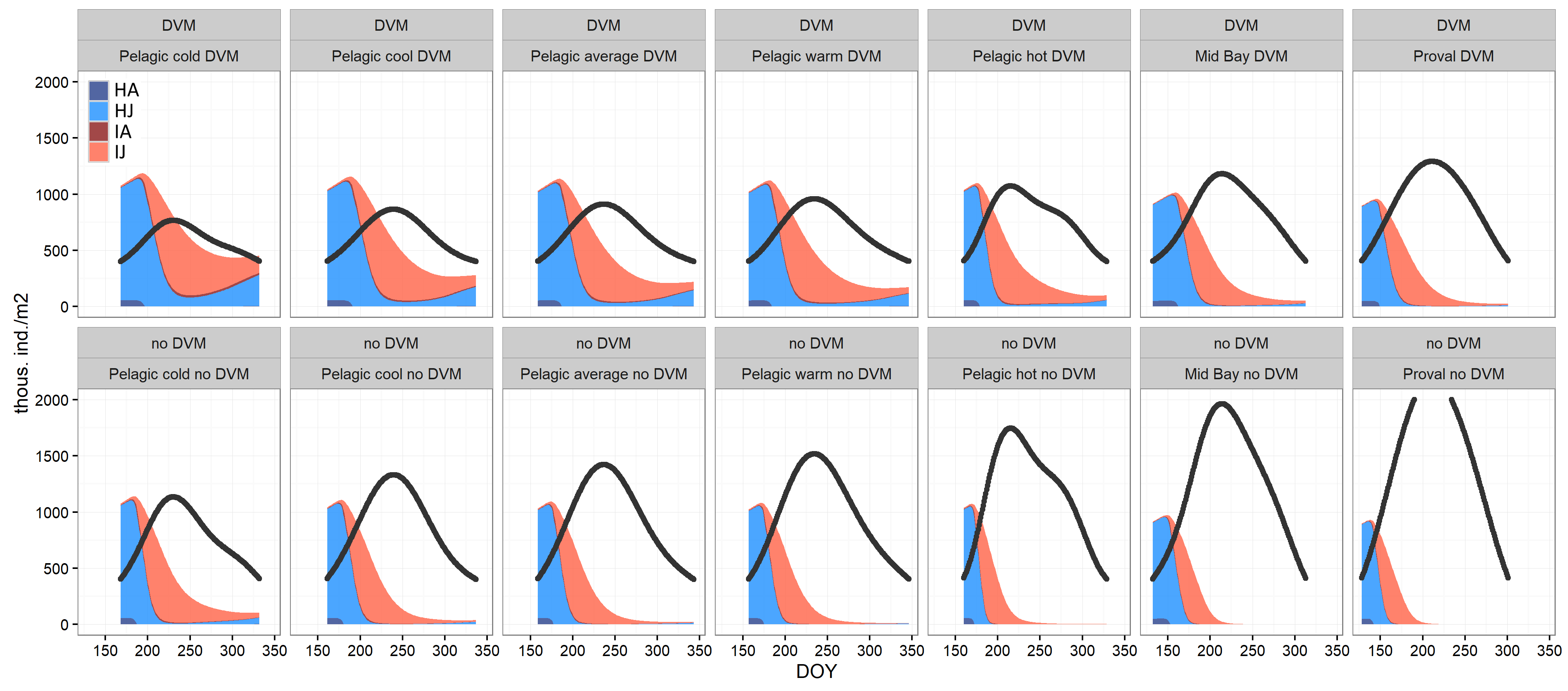

Figure S8: Modeled abundance of *Epischura* during the stratified open water season with temperature-dependent *Saprolegnia* transmission rate (β), modeled based on growth rate of *Saprolegnia* on agar. Runs were performed for 7 different temperature scenarios; each temperature scenario was run for theoretical populations performing DVM and theoretical populations not performing DVM. Filled areas correspond to *Epischura* abundance. HA= healthy adults, HJ= healthy juveniles, IA= infected adults (as thin dark red line below abundance of IJ), IJ= infected juveniles. Black line represents 100*(surface water temperature).
